# Anoxygenic phototrophic *Chloroflexota* member uses a Type I reaction center

**DOI:** 10.1101/2020.07.07.190934

**Authors:** JM Tsuji, NA Shaw, S Nagashima, JJ Venkiteswaran, SL Schiff, T Watanabe, M Fukui, S Hanada, M Tank, JD Neufeld

**Affiliations:** University of Waterloo, 200 University Avenue West, Waterloo, Ontario, Canada, N2L 3G1; Institute of Low Temperature Science, Hokkaido University, Kita-19, Nishi-8, Kita-ku, Sapporo, Japan, 060-0819; Tokyo Metropolitan University, 1-1 Minami-osawa, Hachioji, Tokyo, Japan, 192-0397; Wilfrid Laurier University, 75 University Avenue West, Waterloo, Ontario, Canada, N2L 3C5; Leibniz Institute DSMZ-German Collection of Microorganisms and Cell Cultures GmbH, Inhoffenstrasse 7B, 38124 Braunschweig, Germany

## Abstract

Scientific exploration of phototrophic bacteria over nearly 200 years has revealed large phylogenetic gaps between known phototrophic groups that limit understanding of how phototrophy evolved and diversified. Through Boreal Shield lake water incubations, we cultivated an anoxygenic phototrophic bacterium from a previously unknown order within the *Chloroflexota* phylum that represents a highly novel transition form in the evolution of photosynthesis. Unlike all other known phototrophs, this bacterium uses a Type I reaction center (RCI) for light energy conversion yet belongs to the same bacterial phylum as organisms that use a Type II reaction center (RCII) for phototrophy. Using physiological, phylogenomic, and environmental metatranscriptomic data, we demonstrate active RCI-utilizing metabolism by the strain alongside usage of chlorosomes and bacteriochlorophylls related to those of RCII-utilizing *Chloroflexota* members. Despite using different reaction centers, our phylogenomic data provide strong evidence that RCI- and RCII-utilizing *Chloroflexia* members inherited phototrophy from a most recent common phototrophic ancestor that used RCI, RCII, or both reaction center classes, substantially revising our view of the diversity and evolution of phototrophic life. The *Chloroflexota* phylum preserves an evolutionary record of interaction between RCI and RCII among anoxygenic phototrophs that gives new context for exploring the origins of phototrophy on Earth.

## Main

Chlorophyll-based phototrophy sustains life on Earth through the conversion of light into biologically usable energy^1, 2^. Diverse microorganisms affiliated with at least eight bacterial phyla, discovered over nearly 200 years of scientific exploration^3–11^, perform this key process, and although these bacteria share common phototrophic ancestry^12–14^, many steps in their diversification remain unclear. Substantial gaps in the evolutionary record of phototrophy, apparent through inconsistent topology of photosynthesis gene phylogenies15,16 and a paucity of transition forms between anoxygenic and oxygenic phototrophs^12^, have hindered our ability to answer fundamental questions about the order and timing of phototrophic evolution^15, 17^. Discovery of evolutionary intermediates between known radiations of phototrophic life can help resolve how phototrophs gained their modern functional characteristics.

Anoxygenic phototrophs belonging to the *Chloroflexota* (formerly *Chloroflexi*) phylum have been studied for nearly 50 years^7^, yet their evolutionary properties remain enigmatic. Cultivated from hot spring microbial mats^7, 18, 19^ and mesophilic freshwater environments^20–23^, all known phototrophic *Chloroflexota* representatives use a Type II photosynthetic reaction center (RCII) for light energy conversion^3, 24, 25^. However, several RCII-utilizing phototrophs within the *Chloroflexota* also use chlorosomes, which are bacteriochlorophyll *c*-containing protein:pigment complexes, involved in light harvesting^26, 27^, that are otherwise associated with Type I photosynthetic reaction centers (RCI)^9^. Although structurally homologous, RCI and RCII are thought to have diverged early in the evolution of phototrophy^12–14^ and are well separated in the modern tree of life^3^. For chlorosomes to be present in RCII-utilizing *Chloroflexota* members, some form of genetic interaction, either by descent from a recent common phototrophic ancestor or by lateral gene transfer, must have occurred between RCI- and RCII-utilizing phototrophs during their evolutionary history^28^. Genetic interaction of RCI and RCII has been speculated to have occurred multiple times over the history of life^17^, and is crucial for understanding how oxygenic photosynthesis, the only known process to combine RCI and RCII for electron flow, evolved^6, 12^, yet direct evidence for such genetic interaction has never been reported among anoxygenic phototrophs.

Here we describe the discovery and cultivation of a highly novel member of the *Chloroflexota* phylum that uses RCI, rather than RCII, to perform phototrophy. This phototroph breaks the established paradigm that RCI and RCII are used by phylogenetically distant anoxygenic phototroph clades. Moreover, as a novel transition form in the evolution of photosynthesis, discovery of this phototroph provides strong evidence that RCI- and RCII-utilizing *Chloroflexota* members descended from a most recent common phototrophic ancestor that used RCI, RCII, or both reaction center classes. In this work, we combine physiological, phylogenomic, and environmental metatranscriptomic data to demonstrate RCI-utilizing phototrophic metabolism by a *Chloroflexota* representative, providing fresh context to understand the diversity, origin, and evolution of phototrophic life on Earth.

### Cultivation of a novel Chloroflexota member

With the original intention of cultivating phototrophic *Chlorobia* representatives (phylum *Bacteroidota*), we sampled the illuminated and anoxic water column of an iron-rich Boreal Shield lake (Extended Data Fig. 1a-c) and gradually amended lake water, incubated under light, with a previously published freshwater medium^29^ and ferrous chloride, using Diuron as an inhibitor of oxygenic phototrophs (Extended Data Fig. 1d). Based on 16S ribosomal RNA (rRNA) gene profiles, some of the incubated batch cultures developed high relative abundances of novel microbial populations (inferred from amplicon sequence variants, ASVs) that were only distantly associated with known *Chloroflexota* members (Supplementary Data 1). We used agar-containing medium to further enrich a novel strain, named L227-S17, from one batch culture that represented one of the ASVs from earlier culture profiles (Extended Data Fig. 1e). In addition, from a separate batch culture, we recovered a metagenome-assembled genome (MAG) corresponding to a second novel ASV, named strain L227-5C (Extended Data Fig. 1e; Supplementary Note 1).

After 19 subcultures over four years, strain L227-S17 was brought into a stable enrichment culture that included a putative iron-reducing bacterium, associated with the *Geothrix* genus^30^, named strain L227-G1 (Extended Data Fig. 1f; see additional cultivation details in the Methods section and Supplementary Note 1). Under phototrophic growth conditions, only strains L227-S17 and L227-G1 were detectable in the culture, using 16S rRNA gene amplicon sequencing, to a detection limit of 0.004% (Extended Data Fig. 1f), allowing us to characterize the physiology of strain L227-S17 within a two-member culture system. Based on the RCI-utilizing phototrophic metabolism of L227-S17 (discussed below), we provisionally name the strain “*Candidatus* Chlorohelix allophototropha” (Chlo.ro.he’lix. Gr. Adj. *chloros*, green; Gr. Fem. N. *helix*, spiral; N.L. fem. N. *Chlorohelix*, a green spiral. Al.lo.pho.to.tro’pha. Gr. Adj. *allos*, other, different; Gr. N. *phos*, *-otos*, light; N.L. suff. *-tropha*, feeding; N.L. fem. Adj. *allophototropha*, phototrophic in a different way).

### Phototrophic physiology

We compared the phototrophic properties of the “*Ca.* Chx. allophototropha” enrichment culture to properties of known bacterial phototrophs (Fig. 1; Extended Data Fig. 2). The *in vivo* absorption spectrum of the culture included a strong absorbance peak at 749 nm, which is characteristic of chlorosome-containing phototrophic bacteria^9^ (Fig. 1a and Extended Data Fig. 2a; see Extended Data Table 1 for microbial community data associated with spectroscopy/microscopy-based analyses). Using high-performance liquid chromatography (HPLC), we confirmed that the “*Ca.* Chx. allophototropha” culture contained multiple bacteriochlorophyll *c* species that had absorbance peaks at 435 and 667 nm^9^ (Fig. 1b-c). Large spiralling filaments composed of cells 0.5-0.6 µm wide and 2-10 µm long were visible in the culture (Fig. 1d-e) and were accompanied by smaller rod-shaped cells. The rod-shaped cells corresponded to the *Geothrix* L227-G1 strain based on enrichment of L227-G1 under dark conditions and subsequent microscopy (Extended Data Fig. 2b). Thus, we could establish that “*Ca.* Chx. allophototropha” corresponded to the filamentous cells. The inner membranes of the filamentous cells contained electron-transparent and spherical structures, which matched the expected appearance of chlorosomes after fixation with osmium tetroxide (Fig. 1e; Extended Data Fig. 2c)^31^. Furthermore, “*Ca.* Chx. allophototropha” and the 749 nm absorbance peak were consistently absent when the culture was incubated in the dark (Fig. 1f-g; Extended Data Fig. 2d). Although the culture was typically grown photoheterotrophically to stabilize culture growth, we could also grow the culture photoautotrophically and reproduce the loss of “*Ca.* Chx. allophototropha” in the dark (Extended Data Fig. 2e-f). These data demonstrate that “*Ca.* Chx. allophototropha” L227-S17 is the phototrophic and chlorosome-containing member of the enrichment culture.

**Fig. 1.**
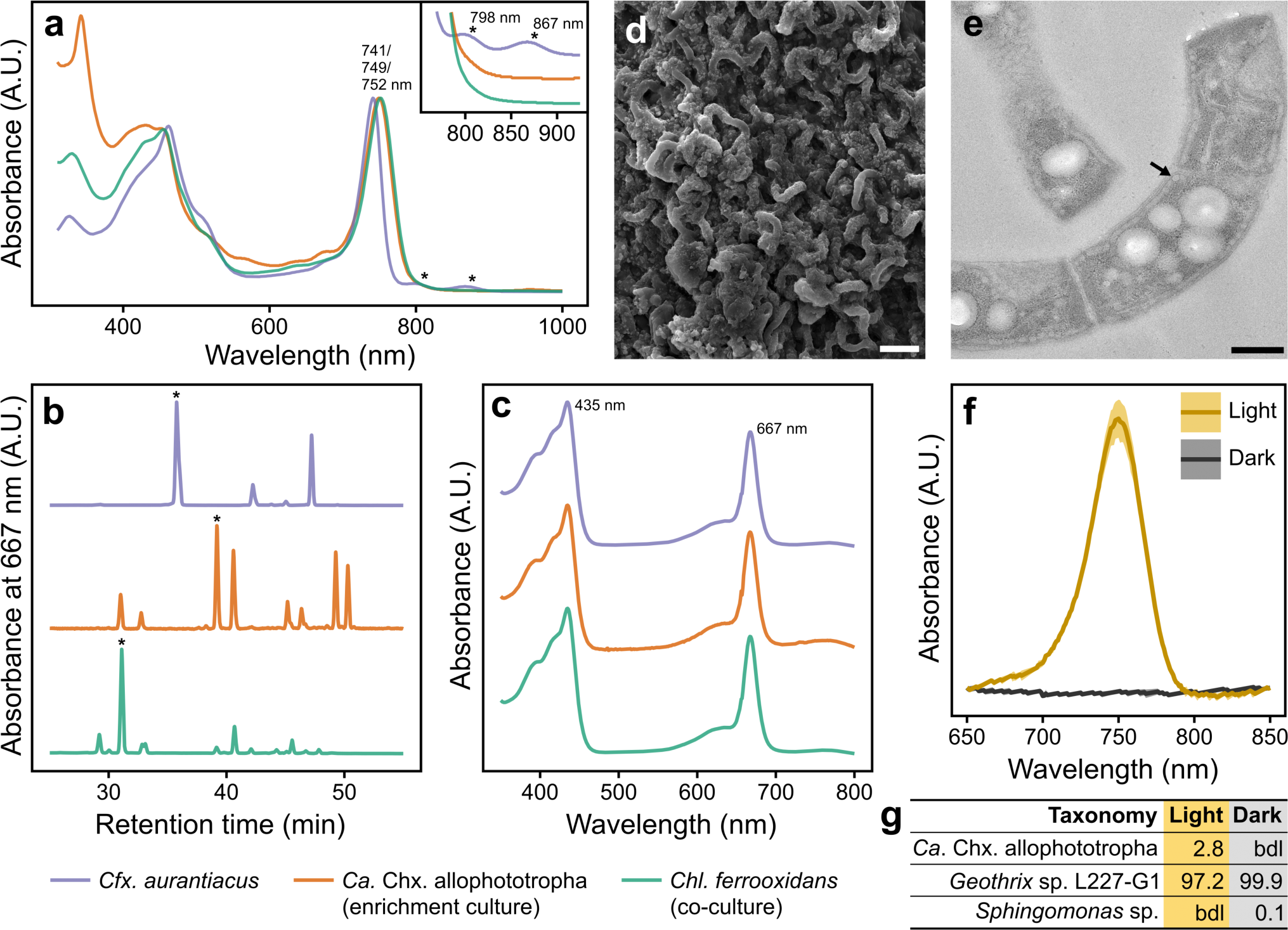
| Phototrophic physiology of the “*Ca.* Chlorohelix allophototropha” culture. **a**, *In vivo* absorption spectrum of the “*Ca.* Chx. allophototropha” culture compared to two reference cultures. The inset shows the ∼760-930 nm region in more detail, with spectra from the three cultures separated on the y-axis. **b-c**, HPLC-based identification of bacteriochlorophyll *c* species in the three cultures. HPLC profiles (**b**) and the *in vivo* absorption spectra associated with the largest HPLC peaks in the profiles (**c**) are shown. Largest HPLC peaks are marked with an asterisk in **b**. All HPLC peaks having >3.5% of the prominence of the largest HPLC peak were associated with the same absorption spectral peaks as shown in **c**. **d**, Scanning electron microscopy image of “*Ca*. Chx. allophototropha” colony material from an early enrichment culture. **e**, Transmission electron microscopy image showing a longitudinal section of “*Ca.* Chx. allophototropha” cells from an early enrichment culture. An example chlorosome-like structure is marked with an arrow. **f-g**, Results of a light vs. dark growth test of the “*Ca.* Chx. allophototropha” culture amended with iron(II) and acetate. *In vivo* absorption spectra (**f**) and relative abundances (in %) of 16S rRNA gene OTUs (**g**) are shown for the culture after two subcultures in the light or dark. Standard deviations (n=3) are shown as shaded areas in **f**; sequencing data in **g** were generated from composite DNA samples. Scale bars in panels **d** and **e** represent 3 μm and 0.3 μm, respectively.

Our spectroscopic data indicate that “*Ca.* Chx. allophototropha” uses a related but novel phototrophic pathway compared to the RCII-utilizing *Chloroflexus aurantiacus*, a chlorosome-containing and phototrophic *Chloroflexota* member that was isolated from a hot spring^7^. Although both strains shared the major chlorosome-associated peak at 740-750 nm in their *in vivo* absorption spectra, the spectrum of *Chloroflexus aurantiacus* had additional absorbance peaks at 798 and 867 nm (Fig. 1a inset). These peaks, as observed previously^32^, may represent the RCII-associated B808-866 complex and were absent in the RCI-associated *Chlorobium ferrooxidans* culture. The peaks were also absent in absorption spectra of suspended whole cells from the “*Ca.* Chx. allophototropha” culture (Extended Data Fig. 2a). The “*Ca.* Chx. allophototropha” and *Chloroflexus aurantiacus* cultures both included multiple bacteriochlorophyll *c* species (Fig. 1b-c), but modifications to bacteriochlorophyll *c* species in the “*Ca.* Chx. allophototropha” culture more closely matched those of the *Chlorobium ferrooxidans* culture (Fig. 1c). Based on these data, the “*Ca.* Chx. allophototropha” culture had RCI-associated physiological properties, despite using chlorosomes and bacteriochlorophyll *c* species like previously known RCII-utilizing *Chloroflexota* members.

### RCI-based phototrophic gene pathway

We identified the RCI-based metabolic potential of “*Ca.* Chx. allophototropha” by analyzing its complete genome sequence. We obtained a single, isolated colony of “*Ca.* Chx. allophototropha” within an agar shake tube (subculture 19.9; Extended Data Fig. 1e) and confirmed that the colony was devoid of the L227-G1 partner strain using 16S rRNA gene amplicon sequencing (Extended Data Table 1). Using hybrid short- and long-read sequencing of DNA from this colony, we assembled the closed “*Ca.* Chx. allophototropha” genome, which consisted of two circular chromosomes (Chr1, 2.96 Mb and Chr2, 2.45 Mb), one circular chromid (375 kb)^33^, and two circular plasmids (241 kb and 55 kb; Extended Data Fig. 3). The chromosomes each encoded at least one copy of identical 16S rRNA genes (Extended Data Fig. 3) and encoded almost entirely (>98%) non-overlapping single-copy marker genes. By similarly performing hybrid short- and long-read DNA sequencing from early enrichments of “*Ca.* Chx. allophototropha” dominated by L227-S17 and L227-G1 (subcultures 15.2 and 15.c, respectively; Extended Data Fig. 1e), we obtained a closed genome bin for the L227-G1 partner strain that consisted of a single circular chromosome (3.73 Mb) and encoded no phototrophic marker genes.

We found no evidence of the RCII-associated *pufLM* genes, used by all known phototrophic *Chloroflexota* members, in the “*Ca.* Chx. allophototropha” genome. Instead, we identified a remote homolog of known RCI genes, analogous to *pscA*^34^, on Chr1 (Fig. 2; Extended Data Fig. 4). The PscA-like primary sequence had only ∼30% amino acid identity to closest-matching RCI sequences in the RefSeq database (as of January 2023), but its predicted tertiary structure included the 11 transmembrane helices^35^ that are expected for a RCI protein, supporting its functional role (Extended Data Fig. 4a). Based on a maximum-likelihood amino acid sequence phylogeny (Fig. 2; Extended Data Fig. 4b), the novel PscA-like predicted protein represents a distinct fifth clade of RCI protein, placing in-between known clades associated with *Chloracidobacteriales*^9^ and *Heliobacteriales*^8^ members. We could replicate our findings of RCI in the MAG of the uncultured L227-5C strain, which encoded a *pscA*-like sequence that placed in the same novel clade (Extended Data Fig. 4b). Both genomes encoded a single copy of the *pscA*-like sequence, suggesting that “*Ca.* Chx. allophototropha” L227-S17 and strain L227-5C use a homodimeric RCI complex for phototrophy^13^. Such distinct phylogenetic placement of the “Ca. Chx. allophototropha” RCI gene excludes the possibility of recent lateral gene transfer from other known phototrophic groups.

**Fig. 2.**
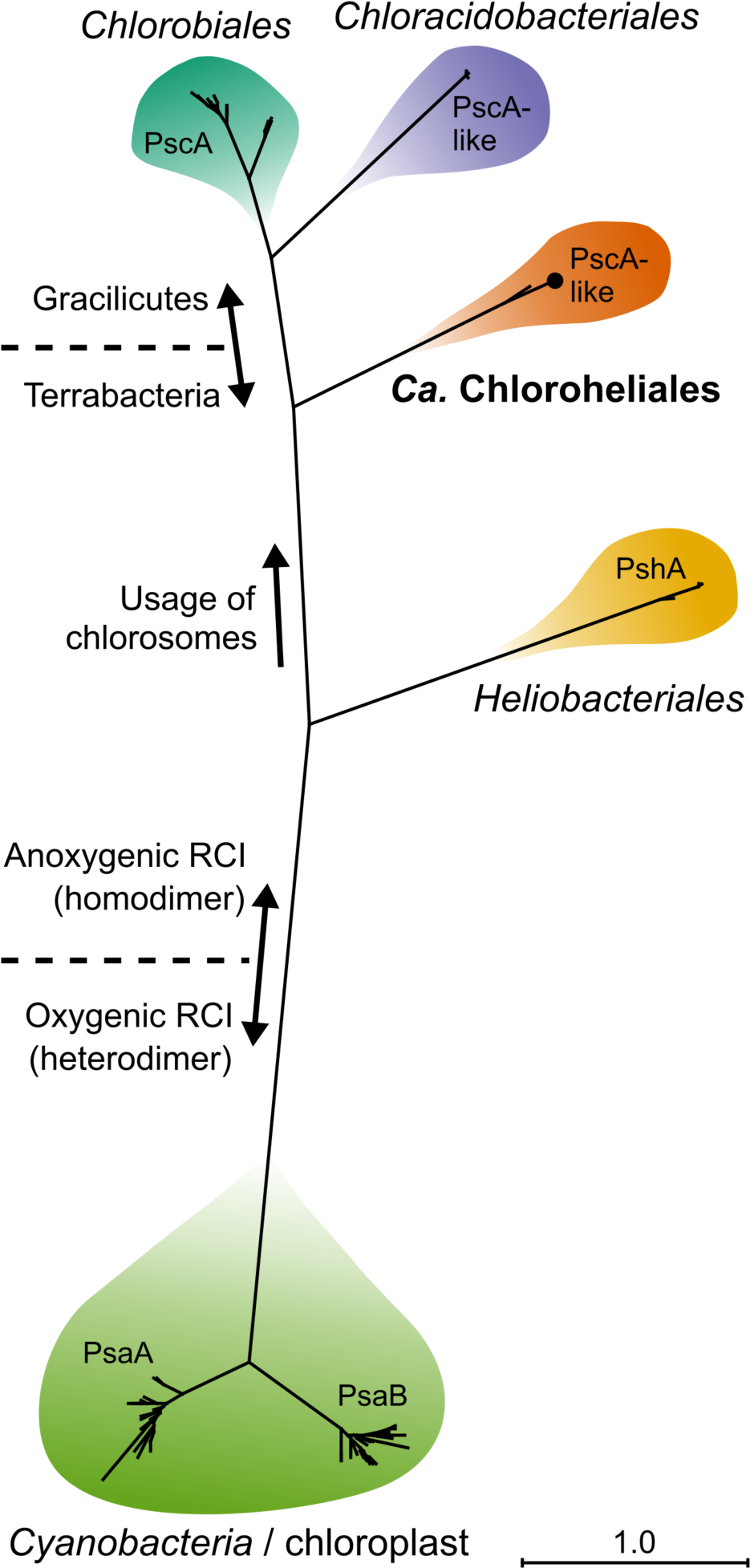
| Type I photosynthetic reaction center of “*Ca.* Chlorohelix allophototropha”. A maximum likelihood phylogeny of RCI primary sequences is shown. Chlorophototrophic lineages are summarized by order name (except *Cyanobacteria* / chloroplasts). The placement of “*Ca*. Chx. allophototropha” is indicated by a black dot. The scale bar shows the expected proportion of amino acid change across the 548-residue masked sequence alignment. All branch points between shaded lineages had 100% bootstrap support.

Along with the *pscA*-like RCI gene, we detected genes for a complete RCI-based phototrophic pathway in the “*Ca.* Chx. allophototropha” genome (Fig. 3; Extended Data Table 2). We detected a homolog of *fmoA*, encoding the Fenna-Matthews-Olson (FMO) protein involved in energy transfer from chlorosomes to RCI^36^, which had 21-26% predicted amino acid identity to known FmoA sequences (Extended Data Fig. 5). This finding makes the *Chloroflexota* the third known phylum to potentially use the FMO protein for phototrophy^9^. The FmoA predicted primary sequence lacked the Cys49 and Cys353 residues previously thought to be necessary for quenching energy transfer^37^. Matching physiological observations, we detected a homolog of the key chlorosome structural gene *csmA*, involved in chlorosome baseplate formation^27, 28^, that had ∼33% identity at the predicted amino acid level to the CsmA primary sequence of *Chloroflexus aurantiacus*^28^. The potential CsmA homolog was missing the His25 previously thought to be involved in bacteriochlorophyll *a* binding^38^. We also detected possible homologs of *csmM* and *csmY*, which encode proteins associated with the chlorosome lipid monolayer, in the “*Ca.* Chx. allophototropha” genome^39^. Such chlorosome-associated proteins are known to be highly diverse and are not necessarily conserved across different phototroph species^27^, meaning that “*Ca.* Chx. allophototropha” may use additional novel proteins in its chlorosomes. In place of Alternative Complex III used by *Chloroflexales* members for photosynthetic electron transport^39^, we found a cytochrome *b_6_f*-like gene cluster that included a putative di-heme cytochrome *c* as described for *Heliobacterium modesticaldum*^40^ (Supplementary Note 2). We detected genes for the entire biosynthesis pathway of bacteriochlorophylls *a* and *c* from protoporphyrin IX, along with a *chlG*-like paralog of *bchG* that may be involved in synthesis of chlorophyll *a* (Extended Data Table 2)^41^. We also identified genes unique to the reductive pentose phosphate (RPP; or Calvin-Benson-Bassham) cycle involved in carbon fixation, including a deep-branching Class IC/ID *rbcL* gene^42^ representing the large subunit of RuBisCO (Extended Data Fig. 6). We did not detect genomic potential for the 3-hydroxypropionate bicycle, which is used for carbon fixation by some RCII-utilizing *Chloroflexales* members, aside from detection of malyl-CoA lyase (MCL), which is also encoded by *Chloroflexota* members incapable of this carbon fixation pathway^43^. At the whole-genome level, we observed no large photosynthetic gene clusters in the “*Ca.* Chx. allophototropha” genome and saw no clear tendency for phototrophy-related genes to be encoded by Chr1 versus Chr2 (Extended Data Fig. 3). Together, genomic data demonstrate that “*Ca.* Chx. allophototropha” has metabolic potential for RCI-driven phototrophy using several highly novel genes compared to known phototrophs.

**Fig. 3.**
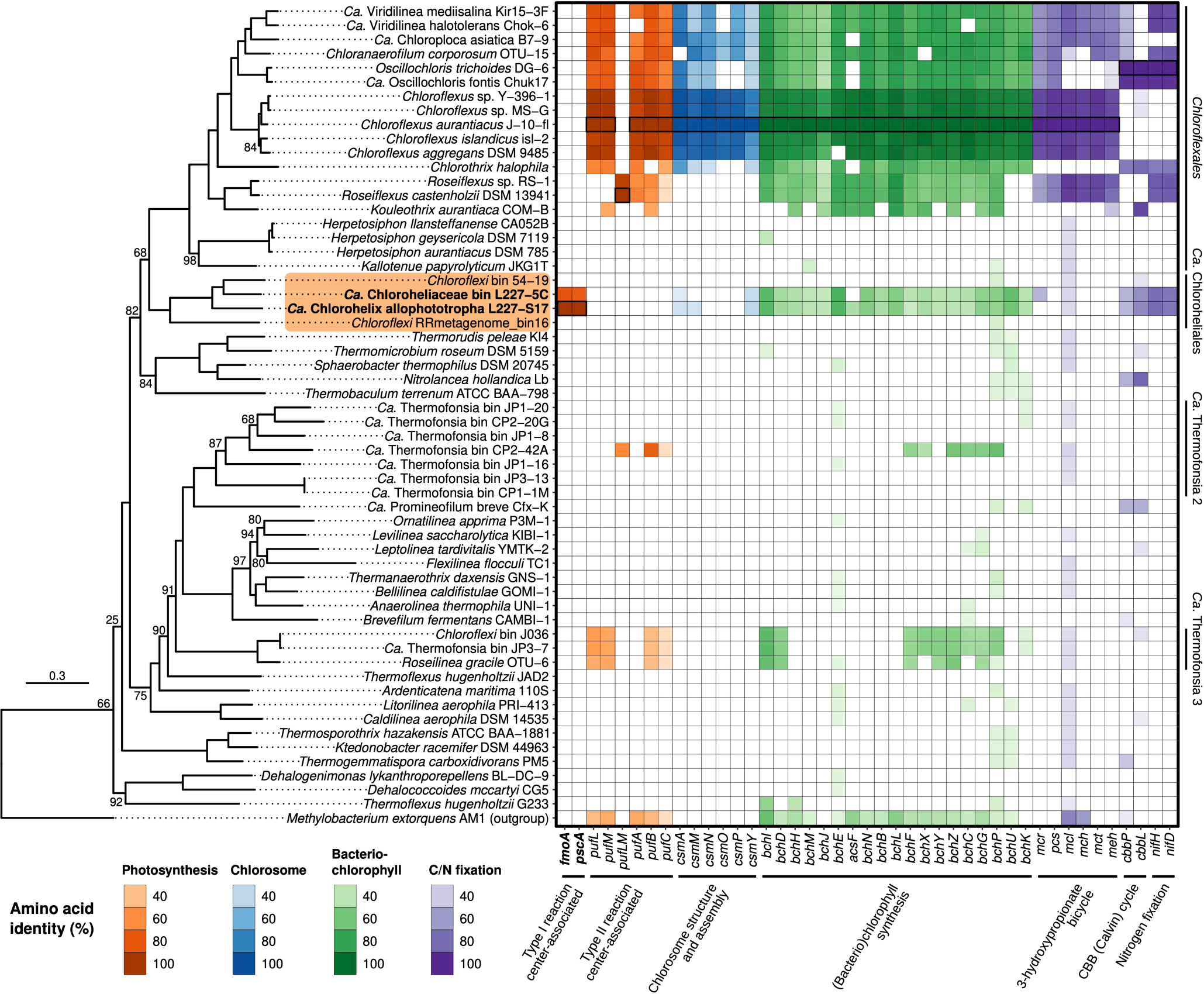
| Genomic potential for phototrophy among members of the *Chloroflexota* phylum. The maximum likelihood phylogeny of *Chloroflexota* members is based on a set of 74 concatenated core bacterial proteins. The scale bar shows the expected proportion of amino acid change across the 11 700-residue masked sequence alignment. Ultrafast bootstrap values (%) are shown for all branches with <100% support. The “*Ca.* Chloroheliales” clade is highlighted in orange, and genomes recovered in this study are in bold. On the right, a heatmap shows the presence/absence of genes involved in photosynthesis or related processes based on bidirectional BLASTP. The fill colour of each tile corresponds to the percent identity of the BLASTP hit. Heatmap tiles of BLASTP query sequences are marked by bolded outlines. Raw results of bidirectional BLASTP are summarized in Supplementary Data 2. Orders containing potential phototrophs are labelled on the right of the heatmap. All genomes in the phylogeny encoded at least 50% of the core 74 proteins except for *Chloroflexi* RRmetagenome_bin16 and “*Ca*. Thermofonsia bin CP2-42A”, which encoded 25 and 34 of the proteins, respectively.

### Detection and activity in Boreal Shield lakes

We examined the ecology and environmental activity of “*Ca.* Chx. allophototropha” relatives in Boreal Shield lakes^44^ to further verify their RCI-based metabolic lifestyle (Fig. 4). Selecting eight seasonally anoxic Boreal Shield lakes nearby (and including) Lake 227, we sampled water columns depth profiles over three years for metagenome and/or metatranscriptome sequencing (Fig. 4a). All but one of the eight lakes developed ferruginous (i.e., iron-rich and sulfate-poor) waters after the onset of anoxia (i.e., all but Lake 626), and the lakes had measurable light penetration into their anoxic zones despite contrasting physicochemical properties, such as dissolved organic carbon and total dissolved iron concentrations (Extended Data Table 3). Searching unassembled metagenome read data, we detected “*Ca.* Chx. allophototropha”-associated *pscA*-like genes in half of the eight seasonally anoxic lakes (i.e., Lakes 221, 304, 222, and 227). Although relative abundances of *pscA*-like gene sequences were low for Lake 227, we detected abundances of up to 1.8% relative to the *rpoB* marker gene in other nearby lakes such as Lakes 221 and 304 (Fig. 4b). Metagenome assembly, genome binning, and bin dereplication allowed us to recover two MAGs from the metagenomes (out of a total of 756 MAGs, having an average completeness and contamination of 82.7% and 2.1%, respectively) that were affiliated with the *Chloroflexota* phylum and encoded a “*Ca.* Chx. allophototropha”-like RCI gene homolog. One of these MAGs, “*Ca*. Chloroheliaceae bin ELA729”, had an average nucleotide identity (ANI) of 99.4% to the MAG of strain L227-5C and likely represents the same species. The second MAG, “*Ca*. Chlorohelix bin ELA319”, had an ANI of 87.5% to the “*Ca.* Chx. allophototropha” genome and could represent a novel but related species. We detected these two dereplicated MAGs in samples from the same four lakes (i.e., Lakes 221, 304, 222, and 227) at >0.01% relative abundance and sometimes across multiple sampling years (Supplementary Data 3), demonstrating that RCI-associated *Chloroflexota* members form robust populations that are widespread among Boreal Shield lakes in this region despite seasonal lake mixing.

**Fig. 4.**
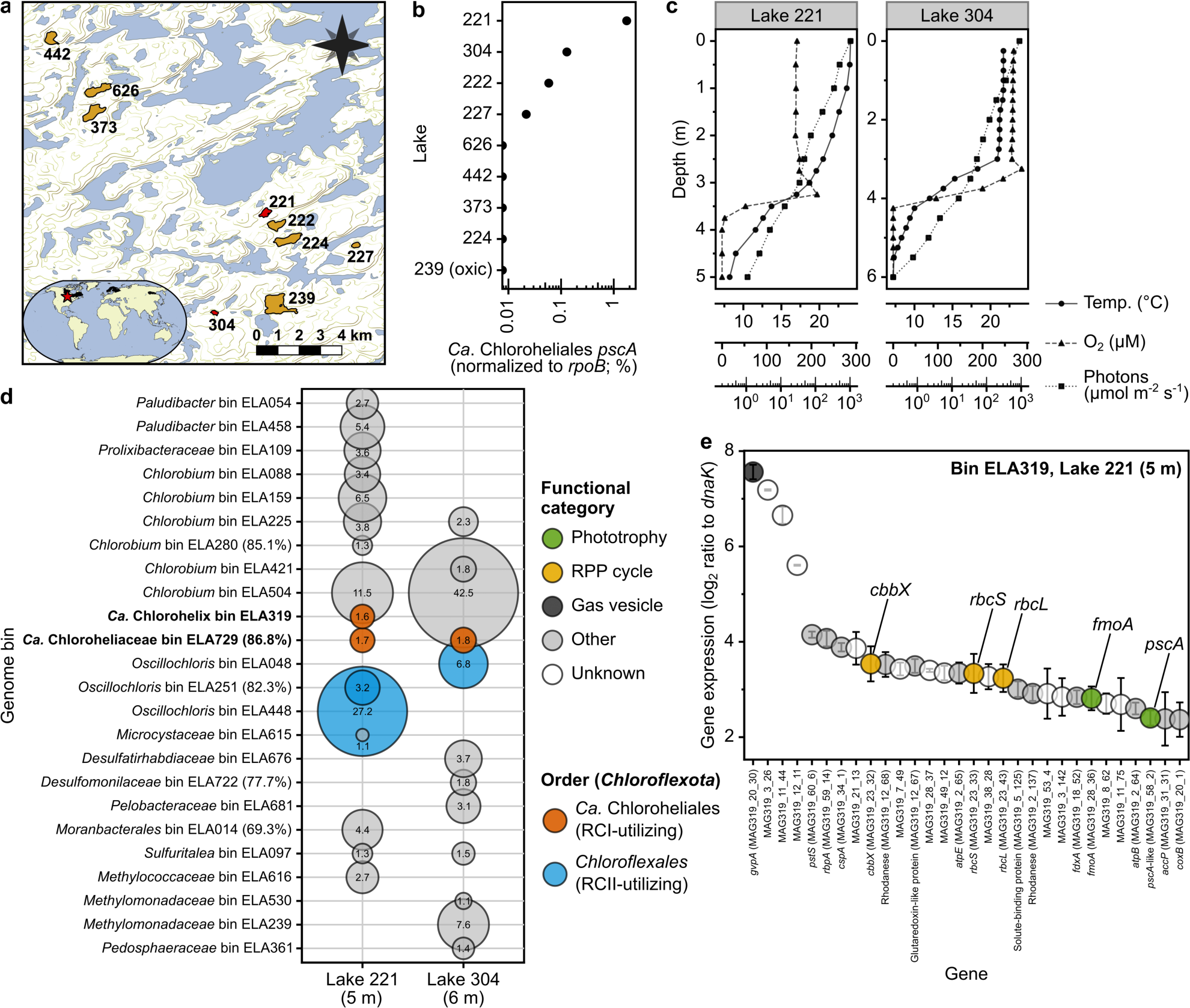
| Distribution and environmental activity of “*Ca.* Chx. allophototropha” relatives in Boreal Shield lakes. **a**, Map showing the nine sampled Boreal Shield lakes at the IISD-ELA. An inset shows the location of the sampling site (red star) and the approximate range of Boreal Shield regions on Earth (black highlights). Spatial data are courtesy of ESRI and DMTI Spatial. **b**, Detection of “*Ca.* Chloroheliales”-like RCI genes in the Boreal Shield lake metagenome dataset. Positive unassembled read hits are shown as a proportion of unassembled reads hits to the housekeeping gene *rpoB*, normalized by gene length. The maximum value is displayed for each lake among all water column metagenomes. **c**, Physical profile data for Lakes 221 and 304, sampled in July 2018. **d**, Bubble plot showing active microbial populations in the water columns of Lakes 221 and 304 based on read mapping of metatranscriptome data (July 2018) to MAGs. The size of each bubble represents the mean relative expression (n=3) of each MAG, as described in the methods, and MAGs having >1% relative expression are shown. (See Supplementary Data 4 for standard deviations, which were on average 7.0% of the mean.) Any MAGs with <90% estimated completion have the completion metric shown beside the MAG ID. **e**, Normalized gene expression of “*Ca.* Chlorohelix bin ELA319” based on Lake 221 metatranscriptome data. The top 30 protein-coding genes with highest normalized expression are shown. Error bars represent the standard deviation of the log_2_ expression ratio (n=3). Highly expressed genes based on Lake 304 metatranscriptome data and read mapping to bin ELA729 are shown in Extended Data Fig. 7, and normalized expression values for all genes are included in Supplementary Data 5.

We sequenced metatranscriptomes from illuminated anoxic water column samples from Lakes 221 and 304 to analyze the *in situ* gene expression of the two MAGs. Like Lake 227 (Extended Data Fig. 1c), Lakes 221 and 304 were shallow lakes with steep chemoclines in the upper 3-4 m of the water column (Fig. 4c). Light penetration into the anoxic zones of these lakes was roughly an order of magnitude higher than Lake 227, where a surface cyanobacterial bloom blocks deep light penetration in summer^44, 45^. Our metatranscriptome data demonstrate that the two RCI-encoding *Chloroflexota* MAGs were highly active compared to other bacterial populations, recruiting as much as 1.8% of mappable metatranscriptome reads from the Lake 221 and 304 samples (Fig. 4d). Both MAGs had upregulated expression of the *pscA*-like RCI gene, the *fmoA* gene, and the *rbcLS* genes encoding RuBisCO, with “*Ca.* Chlorohelix” bin ELA319 (Lake 221) having all of these genes within the top 30 most highly expressed genes in its genome (Fig. 4e; Extended Data Fig. 7). The MAGs also had high expression levels of homologs of *gvpA*, involved in the formation of gas vesicles that may function in buoyancy regulation^46^. Moreover, the MAGs co-occurred with RCII-encoding *Chloroflexota* and RCI-encoding *Chlorobia*-associated MAGs, which were among the highest RNA read-recruiting MAGs in the dataset (Fig. 4d). Our data thus demonstrate that RCI-based phototrophy is actively used by *Chloroflexota* members in natural environments. These RCI-utilizing *Chloroflexota* members potentially form part of a more complex phototrophic microbial consortium in Boreal Shield lake anoxic waters. Given that Boreal Shield lakes number in the millions globally^44^ and might commonly develop iron-rich and anoxic bottom waters, as observed in Fennoscandian lakes geographically distant from the lakes presented in this study^47^, phototrophic consortia including RCI-utilizing *Chloroflexota* members could be relevant to widespread northern ecosystems despite being overlooked previously.

### Phylogenomic properties

“*Ca.* Chx. allophototropha”, the uncultured L227-5C strain, and the two RCI-encoding environmental MAGs belong to a previously uncultivated order within the *Chloroflexota*, based on classification according to the Genome Taxonomy Database (GTDB)^48^. We provisionally name this order the “*Ca.* Chloroheliales” (in place of the former taxon name, “54-19”; Supplementary Note 3). This novel order places within the same class, *Chloroflexia*, as the *Chloroflexales* order that contains RCII-utilizing phototrophs. Based on a concatenated core bacterial protein phylogeny, the “*Ca.* Chloroheliales” order is sibling and basal to the *Chloroflexales* order and is separated from RCII-utilizing phototroph families by the non-phototrophic *Herpetosiphonaceae* family (*Chloroflexales* order)^49^ (Fig. 3). The close phylogenetic placement of these two phototroph-containing orders suggests that RCI- and RCII-utilizing phototrophs have a shared evolutionary history of phototrophy.

We probed the phylogenetic relationships between photosynthesis genes encoded by “*Ca*. Chloroheliales” members and those of other phototrophs to explore their evolutionary relationship (Fig. 5). In a maximum likelihood phylogeny of the chlorosome structural protein CsmA, the “*Ca.* Chloroheliales” clade placed sibling and basal to the RCII-utilizing *Chloroflexota* clade (Fig. 5a). Similarly, in maximum likelihood phylogenies of the (bacterio)chlorophyll synthesis proteins BchIDH/ChlIDH (Fig. 5b) and BchLNB/ChlLNB (Fig. 5c), “*Ca.* Chloroheliales” sequences placed basal to the sister grouping of RCII-utilizing *Chloroflexota* and RCI-utilizing *Bacteroidota* (*Chlorobiales*) members. The placement of the “*Ca.* Chloroheliales” clade was similar in a maximum likelihood phylogeny of BchXYZ proteins (Fig. 5d), except the clade was removed by one branch from placing directly basal to the RCII-utilizing *Chloroflexales* / RCI-utilizing *Bacteroidota* group. Placement of “*Ca.* Chloroheliales” was unstable between BchX, BchY, and BchZ phylogenies, yet in all cases it placed either directly basal to the RCII-utilizing *Chloroflexales* / RCI-utilizing *Bacteroidota* group or placed as the basal member of a clade adjacent to this group (see Fig. 5 caption). Sequences from RCII-utilizing “*Ca.* Thermofonsia” members, which are thought to have acquired phototrophy by recent lateral gene transfer from *Chloroflexales* members^25^, grouped together with sequences of RCII-utilizing *Chloroflexales* members and separately from the novel RCI clade in all phylogenies where these sequences were included. The consistently close phylogenetic placement of phototrophy-associated genes of RCI-utilizing “*Ca.* Chloroheliales” and RCII-utilizing *Chloroflexales* members in all these phylogenies provides strong evidence that, despite using different photosynthetic reaction centers, RCI- and RCII-utilizing *Chloroflexota* members have a shared genetic ancestry of phototrophy.

**Fig. 5.**
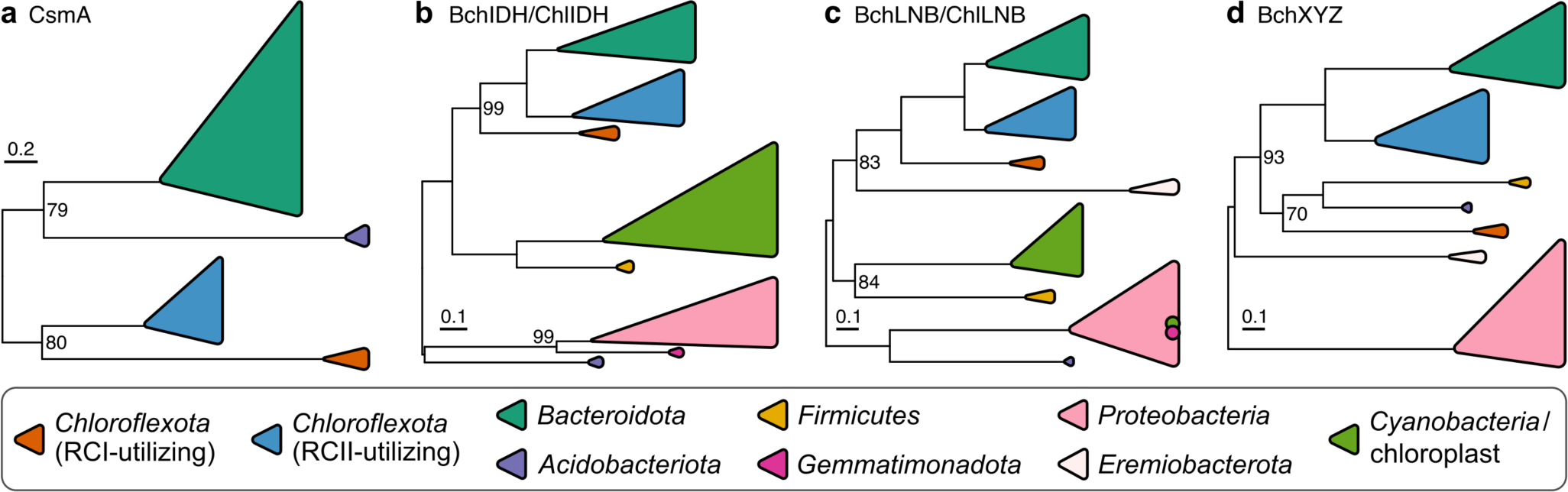
| Phylogenetic relationships of photosynthesis-related genes among known phototrophs. Maximum likelihood phylogenies are shown for the chlorosome baseplate-associated protein CsmA (**a**) and for proteins associated with (bacterio)chlorophyll synthesis: BchIDH/ChlIDH (**b**), BchLNB/ChlLNB (**c**), and BchXYZ (**d**). The phylogenies are midpoint rooted, and ultrafast bootstrap values are shown when <100% between clades. The two dots within the *Proteobacteria* clade (**d**) indicate placement of some *Cyanobacteria* and *Gemmatimonadota* sequences within this clade. Scale bars represent the expected proportion of amino acid change across the masked sequence alignments, which were 74, 1702, 1034, and 988 residues in length for the four panels (**a**, **b**, **c**, **d**), respectively. Detailed versions of the same phylogenies, as well as phylogenies of individual (non-concatenated) genes, are available in the code repository associated with this work.

### Evolutionary implications

Consistent basal phylogenetic placement of “*Ca*. Chx. allophototropha”-like sequences to those of RCII-utilizing *Chloroflexota* members allows robust reconstruction of past genetic interactions between phototrophs in the *Chloroflexota* despite their differing phototrophic properties. Correspondence between photosynthesis gene trees (Fig. 5) and the *Chloroflexota* species tree (Fig. 3) indicates that vertical inheritance, rather than lateral gene transfer, was the dominant factor driving the diversification of RCI- and RCII-utilizing phototrophs within the *Chloroflexia* class. Multiple successive lateral gene transfers would have been needed to explain the phylogenomic patterns in our data, if RCI- and RCII-utilizing *Chloroflexota* members acquired phototrophy genes completely independently. The sister grouping of the *Bacteroidota* (*Chlorobiales*) clade and the clade of RCII-utilizing *Chloroflexota* members in phylogenies of bacteriochlorophyll synthesis genes (Fig. 5b-d) potentially represents a lateral gene transfer event from *Chloroflexota* members to *Bacteroidota* members^28^. Our data therefore provide strong evidence that, despite using different reaction centers, the most recent common ancestor of RCI- and RCII-utilizing *Chloroflexia* members was itself phototrophic, encoding genes for bacteriochlorophyll synthesis and at least one photosynthetic reaction center class.

Two possible models could explain how RCI- and RCII-utilizing *Chloroflexota* members share a recent and common phototrophic ancestor. For the first model, the phototrophic common ancestor utilized only RCI or only RCII. A descendant of that ancestor would have then received genes for the other reaction center via lateral gene transfer while retaining bacteriochlorophyll synthesis genes from the common ancestor. The acquired photosynthetic reaction center would then become functional within that descendant, followed by loss of the original reaction center genes in a “genetic displacement event” as speculated previously^28^. Given that chlorosomes are only associated with RCI outside the *Chloroflexota* and that the “*Ca.* Chloroheliales” places basal to the RCII-associated *Chloroflexota* clade, it is likely that the most recent common ancestor used RCI in this model. For the second model, the common phototrophic ancestor encoded genes for both RCI and RCII. Differential loss of one of the two reaction centers in descendants from this ancestor would then result in the RCI-associated “*Ca.* Chloroheliales” and the RCII-associated *Chloroflexales* orders.

Both the genetic displacement and differential loss models provide an elegant explanation for the presence of chlorosomes among RCII-utilizing *Chloroflexota* members. Chlorosome-associated genes were likely encoded by a phototrophic common ancestor of RCI- and RCII-utilizing *Chloroflexota* members and were then distributed, predominantly by vertical inheritance, to modern RCI- and RCII-utilizing *Chloroflexota* clades. Both models are also relevant to the evolution of oxygenic photosynthesis. In either model, genes for RCI and RCII were likely co-encoded by a single ancestral *Chloroflexota* member at some point during the evolution of the phylum. Rather than integrating RCI and RCII into a single electron transfer pathway, as occurred among the ancestors of oxygenic phototrophs^12^, RCI/RCII gene loss occurred among ancestral *Chloroflexota* members. The *Chloroflexota* phylum may thus record a parallel yet contrasting history of the evolution of phototrophy compared to the *Cyanobacteria*, the only known bacterial phylum to contain oxygenic phototrophs.

Discovery of phototrophs using solely RCI or solely RCII within the same bacterial phylum and with shared photosynthetic ancestry substantially revises our view of the diversity and evolution of phototrophic life (Fig. 6). Our findings represent the first strong evidence that RCI and RCII have interacted outside the *Cyanobacteria*, satisfying previous speculations that such interactions ought to have occurred multiple times over the course of evolution^17^, and provide a rare example of photosynthetic gene movement between physiologically distinct phototroph groups that is necessary for many evolutionary models^6^. In contrast to current paradigms that lateral gene transfer has obscured most signal of the diversification of phototrophy^16^, the distribution patterns of phototrophy-related genes within the *Chloroflexota* can now be understood in light of phylogenomic data from the RCI-encoding transition form “*Ca.* Chx. allophototropha”. Phototrophy within the *Chloroflexota* thus provides a unique case study where enigmatic photosynthesis gene distributions can be resolved through cultivation-based discovery. Moreover, as the only known phylum to contain phototrophs using solely RCI and solely RCII, the *Chloroflexota* represents a new model system to compare the physiological and biochemical bases of RCI- and RCII-driven phototrophy. Discovery of “*Ca*. Chx. allophototropha” will require fundamental views on the diversification of photosynthesis to be revisited in light of this revised view of the phototrophic tree of life.

**Fig. 6.**
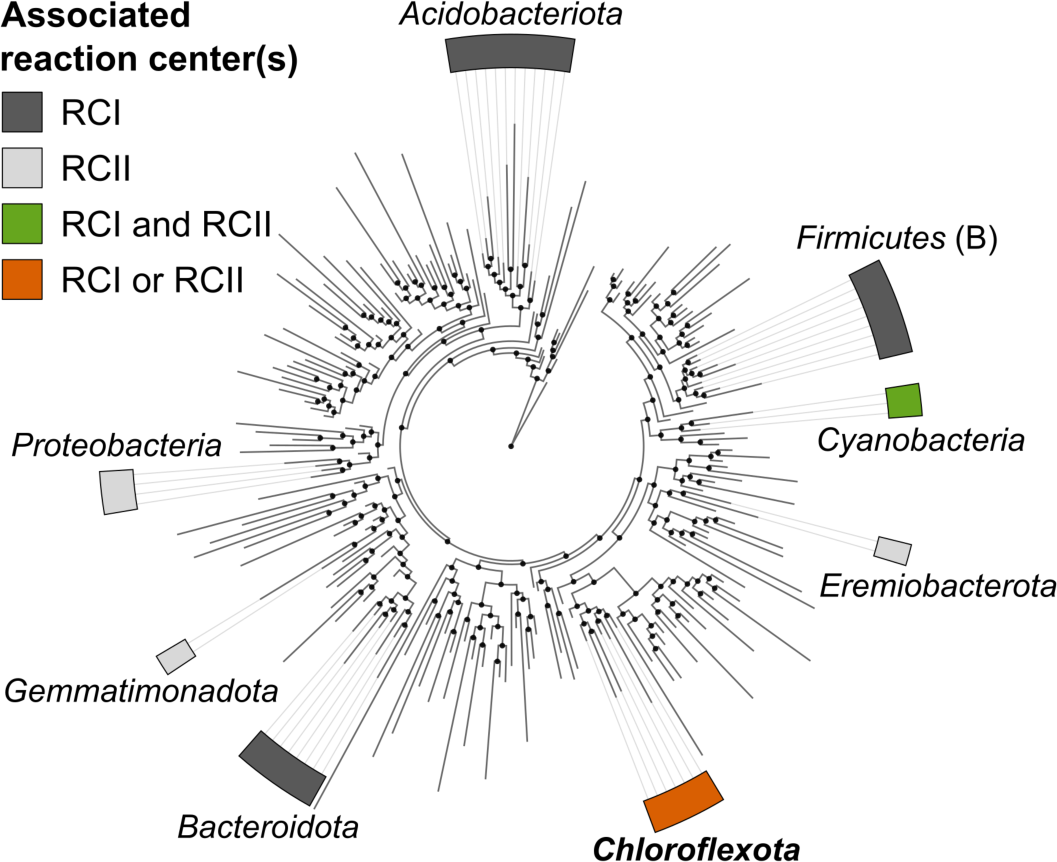
| Revised view of the diversity of phototrophic life. Bacterial phyla with cultured chlorophototrophic representatives are highlighted based on the photosynthetic reaction center class(es) associated with each phylum. The bacterial reference tree from the GTDB (release 89), summarized at the class level by AnnoTree, was used as the phylogeny.

## Acknowledgments

We gratefully acknowledge that our samples were obtained from traditional territory of the Anishinaabe People. We also thank staff at the IISD-ELA and R. Elgood for providing baseline limnological data and sampling advice; J. Mead, E. Barber, J. Wolfe, K. Thompson, E. McQuay, and R. Henderson for assistance with lake sampling; X. Lu, E. Spasov, and L. Shakib, for assistance with DNA and RNA extraction; K. Engel and A. Shinohara for assistance with DNA sequencing; Y. Shirotori and M. Saini for assistance with enrichment cultivation; V. Gaisin for advice for cultivating filamentous phototrophs; H. Ito and R. Tanaka for assistance with pigment analysis; R. Harris and E. Roach for assistance with electron microscopy; T. Chen and Y. Wang for advice about read cloud sequencing; V. Thiel for advice about reference genomes of *Chloroflexota* members; B. Schink for help with taxonomy nomenclature; N. Tran, K. Shimada, Y. Tsukatani, J. Hemp, and R. Tanaka for discussions about biochemistry and genomics; and D. Bryant for critical reading of this work. This work was supported by a Strategic Partnership Grant for Projects from the National Sciences and Engineering Research Council of Canada (NSERC), Discovery Grants from NSERC, a research grant from the Institute for Fermentation, Osaka, a Grant-in-Aid for Scientific Research (20F20384) from the Japan Society for the Promotion of Science (JSPS), and the Grant for Joint Research Program of the Institute of Low Temperature Science, Hokkaido University (20K001). J.M.T. acknowledges a JSPS International Research Fellowship. The work (proposal: 10.46936/10.25585/60000734) conducted by the U.S. Department of Energy Joint Genome Institute (https://ror.org/04xm1d337), a DOE Office of Science User Facility, was supported by the Office of Science of the U.S. Department of Energy operated under Contract No. DE-AC02-05CH11231.

## Author contributions

J.J.V., S.L.S., and J.D.N. conceived the study. J.M.T., N.A.S., and M.T. performed enrichment cultivation. J.M.T. performed pigment analyses and Nanopore DNA sequencing. J.M.T. and N.A.S. performed electron microscopy and read cloud DNA sequencing. J.M.T., J.J.V., and S.L.S. performed the lake survey, and J.M.T. and N.A.S. performed environmental DNA and RNA extraction. J.M.T. analyzed 16S rRNA gene, metagenome, and metatranscriptome sequencing data, conducted phylogenetic and statistical analyses, and visualized the data. J.M.T., S.N., S.H., M.T., and J.D.N. interpreted the data and its impacts on understanding the evolution of photosynthesis. S.H., M.T., T.W., M.F., and J.D.N. supervised research. J.M.T. wrote the paper with N.A.S., S.N., J.D.N., and comments from all other authors.

## Ethics declarations

### Ethics & inclusion statement

All field samples were obtained from the International Institute for Sustainable Development Experimental Lakes Area (IISD-ELA) in northwestern Ontario (Canada) and in partnership with IISD-ELA staff. The IISD-ELA engages and partners with local and regional communities as described at https://www.iisd.org/ela. Researchers affiliated with Canadian institutions led the research project from study design to implementation, own the majority of the data and intellectual property generated from this project, and are co-authors on this work. Field safety training and risk management plans were implemented prior to all field work. Local research is cited as part of this study.

## Methods

### Enrichment cultivation

To culture RCI-utilizing Chloroflexota members, we sampled Lake 227 (49.69° N, 93.69° W), a seasonally anoxic and ferruginous Boreal Shield lake at the International Institute for Sustainable Development Experimental Lakes Area (IISD-ELA). The IISD-ELA sampling site (49.50-49.75° N, 93.50-94.00° W), located near Kenora, Canada, has been described in detail previously^44, 50–53^. Lake 227 develops >100 μM concentrations of dissolved iron in its anoxic water column^44^ (Extended Data Fig. 1c), and anoxia is more pronounced than expected naturally due to the long-term experimental eutrophication of the lake^45^. We collected water from the illuminated portion of the upper anoxic zone of Lake 227 in September 2017, at 3.88 and 5.00 m depth, and transported this water to the laboratory under anoxic and chilled conditions in 120 mL glass serum bottles sealed with black rubber stoppers (Geo-Microbial Technology Company; Ochelata, Oklahoma, USA).

Water was supplemented with 2% v/v of a freshwater medium^29^, amended with 8 mM ferrous chloride, and was distributed anoxically into 120 mL glass serum bottles, sealed with black rubber stoppers (Geo-Microbial Technology Company), that had a headspace of dinitrogen gas at 1.5 atm final pressure. Bottles were spiked with a final concentration of 50 μM Diuron or 3-(3,4-dichlorophenyl)-1,1-dimethylurea (DCMU; Sigma-Aldrich; St. Louis, Missouri, USA) to block oxygenic phototrophic activity^54^. Spiking was performed either at the start of the experiment (for the L227-5C culture, from 5.00 m samples) or as needed following observations of oxygenic phototroph growth (for the L227-S17 culture, from 3.88 m samples). Bottles were incubated at 10°C under white light (20-30 μmol photons m^−2^ s^−1^; blend of incandescent and fluorescent sources), for experiments that resulted in the enrichment of strain L227-5C, or at 22°C under far red LED light (using a PARSource PowerPAR LED bulb; LED Grow Lights Depot; Portland, Oregon, U.S.A.) for experiments that results in the enrichment of strain L227-S17. Cultures were monitored regularly for ferrous iron concentration using the ferrozine assay^55^ and were amended with additional freshwater medium or ferrous chloride when ferrous iron levels dropped, presumably due to iron oxidation (Extended Data Fig. 1d). Up to three subcultures were performed in liquid medium for cultures that depleted ferrous iron (Supplementary Note 1), and the concentration of the liquid freshwater medium was gradually increased during subculturing to 100% (i.e., undiluted) with 8 mM ferrous chloride.

Following initial liquid enrichment, deep agar dilution series^56^ was used to stabilize growth of the “Ca. Chx. allophototropha” culture and to eliminate contaminants. After several rounds of agar cultivation (Supplementary Methods), the agar-containing medium was adjusted to the following composition: 2.80 mM ammonium chloride, 1.01 mM magnesium sulfate, 0.34 mM calcium chloride, 2.20 mM potassium phosphate monobasic, and 11.07 mM sodium bicarbonate, supplemented with 0.5 mL L^−1^ of trace element solution SLA (optionally selenite-free)^57^, a previously published vitamin solution^29^, and selenite-tungstate solution^58^. Concentrations of calcium pantothenate and thiamine were doubled (to 50 and 100 mg L^−1^, respectively) in the vitamin solution compared to the reference. Cobalamin was also added to a final concentration of 25 µg L^−1^, along with the optional addition of resazurin at a final concentration of 0.5 mg L^−1^. The medium was kept at a pH of 7.5 and stored under 90:10 or 80:20 N_2_:CO_2_. We refer to this adjusted freshwater medium as “Chx3.1 medium”. This medium was amended with a final concentration of 0.2-0.8% molten agar, 2 mM ferrous chloride, and 1.2 mM acetate while preparing agar shake tubes, which were then kept stoppered under a N_2_:CO_2_ headspace. Triple-washed Bacto Agar (Becton, Dickinson and Company) or Agar A (Bio Basic; Markham, Canada) could be used as the agar source. (Medium preparation is further described in the Supplementary Methods.) Unless otherwise noted, all “Ca. Chx. allophototropha” agar cultures in subsequent methods were grown using Chx3.1 medium, including 0.2-0.3% (w/v) agar, 2 mM ferrous chloride, and 1.2 mM acetate, and were incubated at 22°C. We also enriched the main contaminating bacterium in the “Ca. Chx. allophototropha” culture, Geothrix sp. L227-G1, as described in the Supplementary Methods.

### Spectroscopic characterization

Cultures of “Ca. Chx. allophototropha” were incubated under 735 nm light via an ISL-150X150 series LED panel (CCS, Kyoto, Japan; ∼30 cm distance at maximum intensity) until green/golden colonies were visible, which were picked and concentrated by centrifugation (Supplementary Methods). A culture of Chlorobium ferrooxidans was grown in Chx3.1 liquid medium with 8 mM ferrous chloride (and no acetate), with medium pH adjusted to ∼6.5, and was incubated at 22°C under white fluorescent light (60 µmol photons m^−2^ s^−1^) until development of brown-coloured iron(III) oxides. Cells were then harvested and concentrated by centrifugation (Supplementary Methods). Lastly, a liquid culture of Chloroflexus aurantiacus J-10-fl was grown in PE medium^18^ under constant illumination (60 W Tungsten lamp, 30 cm illumination distance) at 50°C. For reviving the culture of Chloroflexus aurantiacus from the −80°C glycerol stock, soft agar medium (0.8%) in screw cap test tubes was used. Chloroflexus aurantiacus cells were kept cool during shipping to Hokkaido University and were then harvested by centrifugation (Supplementary Methods). Cell pellets from the three cultures were either analyzed immediately after collection or frozen at −30°C for a maximum of eight weeks before analysis.

To obtain in vivo absorption spectra, culture biomass was sonicated in 10 mM Tris-HCl (pH=8) using a VP-050 ultrasonic homogenizer (TAITEC Corporation; Saitama, Japan). Sonication was performed for 1 min (in ∼10 s/∼10 s on/off pulses), followed by 1 min on ice, for 5 cycles, and resulting crude cell extracts were centrifuged at 10,000 x g for 5 min. Absorption spectra of the supernatant were measured from 300-1000 nm using a UV-1800 UV/Visible Scanning Spectrophotometer (Shimadzu; Kyoto, Japan). Resulting spectra were normalized to have the same peak height at 740-750 nm. For in vitro measurement of bacteriochlorophyll c species, pigments were extracted by gentle disruption of cells diluted at least 1:10 in methanol, followed by centrifugation at 15,000 x g for 10 min and recovery of the supernatant. Supernatant was further diluted in 100% methanol if needed. The extracts were then analyzed by high performance liquid chromatography (HPLC) using a Discovery 5 μm C18 column (Supelco, Sigma-Aldrich). Gradient conditions for the HPLC were as described previously (including a 15 min constant hold of solvent B)^59^. Absorption spectra were measured using a SPD-M10A diode array detector (Shimadzu). Resulting HPLC profiles of absorbance at 667 nm were normalized by maximum peak height. Absorption spectra corresponding to the largest 667 nm peaks in each HPLC profile were also normalized by maximum peak height.

### Light/dark growth test

“Ca. Chx. allophototropha” cultures were incubated for two subculture generations in the light (735 nm, as above) or dark to test the effect on culture growth. Cultures were tested with or without acetate amendment. Triplicate agar shake tubes (that used 0.2% w/v agar in subculture generation 2) were used for all treatments. Cultures were incubated until green/golden colonies were visible in “light” treatments. To harvest biomass from subculture generation 2 tubes, after discarding the top ∼5-10% of the tube contents, soft agar from each tube was centrifuged at 12,000 x g for 5 min at 4°C, followed by removal of the supernatant and an upper agar layer within the pellet. Pellets were washed once in 10 mM Tris-HCl (pH=8) and frozen at −30°C for microbial community analysis. After DNA extraction (described below), DNA extracts for replicate samples were combined in equal DNA mass ratios for 16S rRNA gene sequencing. For acetate treatments, partial in vivo spectra were generated from a portion of the unfrozen pellet. Sonication was performed as described for in vivo spectra above, except two sonication cycles were used instead of five. Crude cell extracts were then centrifuged at 5000 x g for 1 min at 4°C, and the absorption spectrum of the supernatant was recorded from 500 to 1000 nm, to capture the chlorosome-associated peak, using a UV-1800 UV/Visible Scanning Spectrophotometer (Shimadzu). Resulting absorption spectra were normalized using linear baseline correction between 650-850 nm.

### Electron microscopy

To perform transmission electron microscopy (TEM), cell biomass was picked from “Ca. Chx. allophototropha” cultures grown under white light (30 µmol photons m^−2^ s^−1^; mix of incandescent and fluorescent sources), and residual agar surrounding cells was digested using agarase. One unit of β-agarase I (New England Biolabs; Ipswich, Massachusetts, USA) and 10 µL of 10x reaction buffer was added to 100 μL of cell suspension and incubated at 42°C for 1.5 h. Following cell pelleting and removal of supernatant, cells were then fixed for 2 h at 4°C in 4%/4% glutaraldehyde/paraformaldehyde (dissolved in phosphate-buffered saline) and stored at 4°C. Sample preparation, including fixation with osmium tetroxide, and imaging was performed at the Molecular and Cellular Imaging Facility of the Advanced Analysis Center (University of Guelph, Ontario, Canada), as described in the Supplementary Methods. For scanning electron microscopy (SEM), cultures were grown as above but also included 120 µM sulfide. Fixed cells, digested with agarase as above, were prepared and imaged at the Molecular and Cellular Imaging Facility of the Advanced Analysis Center (University of Guelph; Supplementary Methods).

### 16S rRNA gene amplicon sequencing and analysis

To confirm the microbial community composition of enrichment cultures, genomic DNA was extracted from pelleted cell biomass using the DNeasy UltraClean Microbial Kit (Qiagen; Venlo, The Netherlands). For early enrichment cultures (L227-S17 and L227-5C subculture generations 0 to 1; Supplementary Data 1), a 10 min treatment at 70°C was performed after adding Solution SL to enhance cell lysis. Resulting DNA extracts were quantified using the Qubit dsDNA HS Assay Kit (Thermo Fisher Scientific; Waltham, Massachusetts, U.S.A.) Three different 16S rRNA gene amplicon sequencing methods were then used to analyze enrichment culture samples (Extended Data Table 1). For some cultures, the V4-V5 region of the 16S rRNA gene was amplified from extracted DNA via the universal prokaryotic PCR primers 515F-Y^60^ and 926R^61^ as described previously^62–64^. Library pooling, cleanup, and sequencing on a MiSeq System (Illumina; San Diego, California, USA) was performed as described previously^62^ to generate 2×250 bp paired-end reads. For other cultures, the V4 region was amplified and sequenced by the Bioengineering Lab Co. (Sagamihara, Japan) using the universal prokaryotic primers 515F^65^ and 806R^65^. Sequencing was performed on a MiSeq System to generate 2×300 bp paired-end reads. For a final set of cultures, the 16S Barcoding Kit 1-24 (Oxford Nanopore Technologies; Oxford, U.K.) was used to amplify the near full-length 16S rRNA gene (V1-V9 region) using the universal bacterial PCR primers 27F^66^ and 1492R^66^. Resulting libraries were sequenced on a R9.4.1 Flongle flow cell (FLO-FLG001; Oxford Nanopore Technologies), and basecalling was performed using Guppy version 5.0.16 or 5.1.12 (Oxford Nanopore Technologies) via the Super Accuracy model.

Sequence data analysis was performed for V4-V5 region samples using QIIME2^67^ version 2019.10 via the AXIOME3^68^ pipeline, commit 1ec1ea6 (https://github.com/neufeld/axiome3), with default parameters. Briefly, paired-end reads were trimmed, merged, and denoised using DADA2^69^ to generate an ASV table. Taxonomic classification of ASVs was performed using QIIME2’s naive Bayes classifier^70^ trained against the SILVA SSU database^71, 72^, release 132. The classifier training file was prepared using QIIME2 version 2019.7. For V4 region samples, QIIME2 version 2022.8 was used to analyze the samples directly. After removing forward and reverse PCR primers from the reads using CutAdapt^73^, reads were trimmed on the 3’ ends, merged, and denoised using DADA2^69^ to generate an ASV table. For V1-V9 region samples, NanoCLUST commit a09991c (fork: https://github.com/jmtsuji/nanoclust) was used to generate polished 16S rRNA gene sequence clusters^74^, followed by primer trimming, 99% OTU clustering, and chimera removal as described in the Supplementary Methods. Resulting amplicon sequences (ASVs or OTUs) were then classified as “Ca. Chx. allophototropha” or Geothrix sp. L227-G1 based on a 100% match across their complete sequence to reference 16S rRNA gene sequences generated during genome sequencing for these species. For one deeply sequenced V4 region sample (subculture 21.2; Extended Data Fig. 1f), a 1-base mismatch to the “Ca. Chx. allophototropha” or Geothrix sp. L227-G1 sequence was allowed during classification to assign taxonomy to four low count ASVs (<0.3% relative abundance each) that may represent sequencing artefacts.

### Metagenome sequencing of enrichment cultures

The functional gene content of early liquid enrichment cultures (subculture generations 0-1) was assessed via shotgun metagenome sequencing. Genomic DNA was extracted as above, and library preparation and sequencing were performed at The Centre for Applied Genomics (TCAG; The Hospital for Sick Children, Toronto, Canada). The Nextera DNA Flex Library Prep Kit (Illumina) kit was used for metagenome library preparation, and libraries were sequenced using a HiSeq 2500 System (Illumina) with 2×125 base paired-end reads to obtain 5.0-7.3 million reads pairs per sample.

To close the genome of “Ca. Chx. allophototropha”, a single large colony was picked from an agar shake tube of subculture 19.9, which was grown with 0.4% (w/v) agar under 735 nm light (as for in vivo spectra above) and included an additional 100 µM sulfide. Genomic DNA was extracted from the picked colony as above, and short read metagenome sequencing was then performed by the Bioengineering Lab Co. (Sagamihara, Japan). Briefly, input DNA was processed using the Nextera XT DNA Library Prep Kit (Illumina) followed by the MGIEasy Circularization Kit (MGI; Shenzhen, China), and the resulting library was sequenced as 2×200 bp paired-end reads on a DNBSEQ-G400 (MGI) to generate 3.3 million reads pairs. Remaining material from the same DNA extract of subculture 19.9 was then concentrated using the DNA Clean and Concentrator-5 kit (Zymo Research; Irvine, California, U.S.A.) and used for long read sequencing. Using 225 ng of the DNA sample, spiked with 150 ng of Lambda DNA (EXP-CTL001; Oxford Nanopore Technologies), a long read sequencing library was prepared using the Ligation Sequencing Kit (SQK-LSK110; Oxford Nanopore Technologies) with Long Fragment Buffer, followed by sequencing using a R9.4.1 Flongle flow cell (FLO-FLG001; Oxford Nanopore Technologies). Reads matching the Lambda phage genome (NC_001416.1) were depleted during sequencing using adaptive sampling. Basecalling was performed using Guppy 5.0.16 (Oxford Nanopore Technologies) with the Super Accuracy model, generating 0.51 million reads with a mean length of 2.1 kb.

In addition, to close the Geothrix L227-G1 genome bin, we performed short and long-read DNA sequencing on two enrichment cultures of “Ca. Chx. allophototropha” (subcultures 15.2 and 15.c) as described in the Supplementary Methods.

### Enrichment culture metagenome assembly and genome binning

Short read metagenome sequencing data from early enrichment cultures (subcultures 0-1) and from “Ca. Chx. allophototropha” subculture 15.2 were analyzed using the ATLAS pipeline, version 2.2.0^75^, to generate a set of dereplicated MAGs (Supplementary Methods). To assemble the complete “Ca. Chx. allophototropha” genome, we used hybrid long- and short-read sequencing data from subculture 19.9. Nextera sequencing adapters on the 5’ and 3’ ends of short reads were trimmed using CutAdapt version 3.4^73^. Short read QC was then performed using the ‘qc’ module of ATLAS version 2.8.2^75^. Within ATLAS, sequencing adapters used during MGI-based short read library preparation were added to the ‘adapters.fa’ file to facilitate adapter removal. The QC-processed short read data was then combined with long read data to perform hybrid genome assembly using a custom pipeline we developed named Rotary (https://github.com/jmtsuji/rotary, doi:10.5281/zenodo.6951912), commit e636236. Within Rotary, long read QC was performed using BBMap version 37.99 (Bushnell B. – <u>sourceforge.net/projects/bbmap/</u>) to remove reads shorter than 1 kb or with quality score <13. The long reads were then assembled using Flye version 2.9-b1768 with the ‘--nano-hq’ input flag in ‘meta’ mode^76^. Potential short gaps or overlaps on the ends of circular contigs were corrected using a customized wrapper of the ‘merge’ module of Circlator version 1.5.5^77^. Assembled contigs were then polished using Medaka 1.4.4 (Oxford Nanopore Technologies) with the ‘r941_min_sup_g507’ model, and short reads were mapped to polished contigs using BWA-MEM version 0.7.17^78^. Short read polishing was then performed on contigs using Polypolish version 0.5.0^79^, followed by a second round of short read polishing via POLCA^80^ version 4.0.8. After short read polishing, only contigs with a short read coverage depth of >10x were retained for downstream analysis. Circular contigs were then rotated using the ‘fixstart’ module of Circlator version 1.5.5. The dnaA gene, identified using the profile hidden Markov model (HMM) ‘Bac_DnaA’ (PF00308.21, Pfam^81^) via hmmsearch version 3.3.2^82^, was set as the start point of the applicable contigs, based on coding sequence predictions from Prodigal version 2.6.3^83^. A final round of short read polishing, using Polypolish as above, was then performed on rotated circular contigs. After running Rotary, single copy marker genes were identified in the genome using CheckM version 1.0.18^84^, and the genome was annotated using PGAP version 2022-02-10.build5872^85^. The closed genome bin of the Geothrix L227-G1 strain was similarly assembled from short- and long-read DNA sequencing data from subcultures 15.2 and 15.c, followed by genome binning (Supplementary Methods). We mapped reads from all metagenomes against a dereplicated set of genomes and MAGs resulting from these analyses to compare the microbial community composition of the enrichment cultures (Supplementary Methods).

### Identification of RCI-associated genes

We searched for RCI-associated gene homologs in the genomes of strains L227-S17 and L227-5C using hmmsearch^82^ version 3.1b2 and HMMs downloaded from Pfam^81^. Genes encoding the core Type I reaction center (pscA/pshA/psaAB; PF00223), a Type I reaction center-associated protein (Chlorobia-associated pscD; PF10657), chlorosomes structural units (csmAC; PF02043, PF11098), and a bacteriochlorophyll a binding protein (fmoA; PF02327) were queried. We also queried genes encoding an RCI-associated iron-sulfur protein (pscB; PF12838) and RCI-associated c-type cytochromes (Chlorobia-associated pscC or Heliobacterium-associated PetJ; PF10643 or PF13442), although HMMs used for these genes were non-specific. The genomes were confirmed to lack the pufLM genes associated with RCII using the “Photo_RC” HMM (PF00124), which targets both the pufL and pufM genes associated with the RCII core reaction center. We used the same HMM set (excluding non-specific HMMs mentioned above) with an e-value threshold of 10^−1^ to confirm the lack of photosynthesis-associated marker genes in the Geothrix sp. L227-G1 genome.

The tertiary structures of the detected pscA-like gene homologs in the genomes of “Ca. Chx. allophototropha” L227-S17 and strain L227-5C were predicted using I-TASSER^86^. Custom HMMs were also built for the pscA-like gene and the fmoA gene homologs encoded by the strains. Primary sequences were aligned using Clustal Omega^87^ version 1.2.3, and HMMs were generated using hmmbuild^82^ version 3.1b2. Custom HMMs and homology models generated by I-TASSER are available in the code repository associated with this work.

### Assessment of genomic potential for photosynthesis within the Chloroflexota phylum

We collected representative genomes from Chloroflexota phylum members (Supplementary Methods) and used GToTree^88^ version 1.4.11 and IQ-TREE^89^ version 1.6.9 to construct a species tree. Within GToTree, the ‘Bacteria.hmm’ collection of 74 single-copy marker genes was used to generate a concatenated protein sequence alignment, with a minimum threshold of 30% of marker genes per genome. All but two genomes contained >50% of the marker genes (Fig. 3 caption). A maximum likelihood phylogeny was built by IQ-TREE using the masked multiple sequence alignment, with the LG+F+R6 evolutionary model as determined by ModelFinder^90^ and 1000 ultrafast bootstraps^91^.

A collection of photosynthesis-associated genes, including genes associated with photosynthetic reaction centers, antenna proteins, chlorosome structure and attachment, bacteriochlorophyll synthesis, and carbon fixation, was selected based on the genome analyses of Tang and colleagues^39^ and Bryant and colleagues^28^. Reference sequences for these genes were selected from genomes of well-studied representatives of the Chloroflexota phylum, namely Chloroflexus aurantiacus^7^, Oscillochloris trichoides^20^, and Roseiflexus castenholzii^92^. Bidirectional BLASTP^93^ was performed on reference sequences against the entire Chloroflexota genome collection to detect potential orthologs. The BackBLAST pipeline^94^, version 2.0.0-alpha3 (doi:10.5281/zenodo.3697265), was used for bidirectional BLASTP, and cutoffs for the e value, percent identity, and query coverage of hits were empirically optimized to 10^−3^, 20%, and 50%, respectively. Outside the bidirectional BLASTP search, we identified additional genes potentially involved in the bacteriochlorophyll synthesis pathway in the “Ca. Chx. allophototropha” genome based on ^41^, and we identified genes associated with photosynthetic electron transport as described in Supplementary Note 2.

### Phylogenetic assessment of photosynthesis-associated genes

We compared genes associated with RCI/PSI (pscA/pshA/psaAB)^95^, bacteriochlorophyll a binding (fmoA)^36^, chlorosome structure (csmA)^38^, (bacterio)chlorophyll synthesis (bchIDH/chlIDH, bchLNB/chlLNB, and bchXYZ)^28^, and carbon fixation via the RPP cycle (rbcL)^96^ between “Ca. Chloroheliales” members and other known phototrophs. Genes were identified among a reference genome set as described in the Supplementary Methods. Predicted primary sequences were aligned using Clustal Omega^87^ version 1.2.3, followed by manually inspection. Alignments were masked using Gblocks^97^ version 0.91b with relaxed settings (-t=p -b3=40 -b4=4 -b5=h) to preserve regions with remote homology. Maximum likelihood protein phylogenies were then built using IQ-TREE^89^ version 1.6.9 with 1000 rapid bootstraps to calculate branch support values^91^. Evolutionary rate models, identified using ModelFinder^90^, were as follows: LG+F+G4 (RCI/PSI), LG+G4 (FmoA), LG+F+G4 (CsmA), LG+F+I+G4 (BchIDH/ChlIDH), LG+F+I+G4 (BchLNB/ChlLNB), LG+I+G4 (BchXYZ), and LG+I+G4 (RbcL).

### Boreal Shield lake survey

We sampled eight seasonally anoxic lakes (Lakes 221, 222, 224, 227, 304, 373, 442, and 626) within the IISD-ELA, along with a permanently oxic reference lake (Lake 239). The water columns of the lakes were sampled in the summer or fall of 2016-2018 across four main sampling events (Supplementary Methods). During lake water column sampling, temperature and dissolved oxygen were measured using an EXO multi-parameter sonde (Xylem; Rye Brook, New York, U.S.A.), although for selected sampling events a HQ40D Portable Multi Meter for Water (Hach; Loveland, Colorado, U.S.A.) or handheld water quality meter (Xylem) were used. Light intensity was measured using a LI-192 underwater quantum sensor (LI-COR Biosciences; Lincoln, New England, U.S.A.). Total dissolved iron was measured using the ferrozine assay^55, 98^ on water samples that were filtered in-line (via 0.45 µm membrane filters) and preserved in 0.5 N HCl, and sulfate and dissolved organic carbon samples were collected and measured as described previously^44^.

Samples for water column DNA were collected by pumping water, using a closed system gear pump and line, through sterile 0.22 µm Sterivex polyvinyl fluoride filters (Merck Millipore; Burlington, Massachusetts, U.S.A.). Water column RNA samples were collected similarly, except filter cartridges were immediately filled with 1.8 mL of DNA/RNA Shield (Zymo Research) once packed and purged of residual water. Filters were collected (and subsequently extracted and analyzed) in triplicate for RNA. Sterivex filters were kept chilled after collection until being frozen (at −20°C) upon return to the sampling camp the same day. Filters were then shipped chilled to the University of Waterloo and were kept frozen (at −20°C) until processing.

### Environmental DNA/RNA extraction and sequencing

Environmental DNA was extracted from the excised membranes of Sterivex filters using the DNeasy PowerSoil or DNeasy PowerSoil HTP 96 Kit (Qiagen; Venlo, The Netherlands). Environmental RNA extraction was performed using the ZymoBIOMICS DNA/RNA Miniprep Kit (Zymo Research) via the “DNA & RNA Parallel Purification” protocol with in-column DNase I treatment. Modifications to the standard kit protocols for use of Sterivex filters are described in the Supplementary Methods.

Metagenome sequencing was performed for June and September 2016 samples by the U.S. Department of Energy Joint Genome Institute (Lawrence Berkeley National Laboratory). The Nextera XT DNA Library Preparation Kit (Illumina; including library amplification steps) was used, followed by 2×150 bp paired-end read sequencing, using a HiSeq 2500 System, to generate 44.1 to 136.8 million read pairs per sample. Metagenome sequencing for 2017 and 2018 field samples was performed by the McMaster Genome Facility (McMaster University; Hamilton, Ontario, Canada). Sequencing libraries were constructed using the NEBNext Ultra II DNA Library Prep Kit for Illumina (New England Biolabs), including library amplification steps, using input DNA sheared via an ultrasonicator (Covaris; Woburn, Massachusetts, U.S.A.). The resulting library was sequenced using a HiSeq 2500 System with 2×200 bp paired-end reads, followed by a second sequencing run using a HiSeq 2500 System with 2×250 bp paired-end reads. Reverse reads for the 2×200 bp run were truncated at 114 bp due to a sequencer error. After pooling of data from both runs, a total of 20.6 to 64.1 million read pairs were generated per sample. Metatranscriptome sequencing for 2017 and 2018 field sampling was also performed by the McMaster Genome Facility (McMaster University). Following rRNA depletion using the Ribo-Zero rRNA Removal Kit (Bacteria; Illumina), library preparation was performed using the NEBNext Ultra II RNA Library Prep Kit for Illumina (New England Biolabs) using normalized RNA inputs per sample and without direction RNA selection. The resulting library was sequenced on a portion of a lane of a HiSeq 2500 System in Rapid Run Mode with 2×200 bp paired-end reads. This was the same sequencing run as for 2017-2018 metagenomes. Sequencing generated 5.5 to 10.2 million read pairs per replicate. In total, 37 metagenomes were sequenced, along with 9 metatranscriptomes representing 3 samples (see Extended Data Table 3 and Supplementary Data 3-4).

### Metagenome and metatranscriptome analysis

All environmental metagenome data was processed using the ATLAS pipeline, version 2.1.4^75^. Default settings were used except that the minimum percent identity threshold for read mapping (via ‘contig_min_id’) was set to 99%, and only MaxBin2 and MetaBAT2 were used as binning algorithms^99, 100^. To enhance genome binning quality, six lake metagenomes that were previously sequenced from the water columns of Lakes 227 and 442^101^ were included in the ATLAS run, along with a single metagenome from the aphotic zone of the nearby and meromictic Lake 111, which was sampled in July 2018. The Lake 111 metagenome was not analyzed further in the context of this work. The entire ATLAS pipeline, including quality control on raw reads, metagenome assembly of individual samples, metagenome binning, dereplication of bins from all samples and bin analysis, and gene clustering and annotation, was run end-to-end. For the genome binning step, all metagenome samples from the same lake were summarized in the same ‘BinGroup’, allowing for differential abundance information between samples from the same lakes to be used to guide genome binning. After running ATLAS, dereplicated MAGs were taxonomically classified using the GTDB-Tk, version 0.3.2^102^, which relied on the GTDB, release 89^48^. All MAGs had a minimum percent completeness of 50% and maximum percent contamination of 10% based on CheckM^84^. The relative abundance of each MAG in a sample was calculated by dividing the number of QC-processed reads mapped to the MAG by the total number of assembled reads (i.e., raw reads that mapped to assembled contigs) for that sample.

Metagenomes were also searched at the unassembled read level for “Ca. Chx. allophototropha”-associated pscA-like genes. Short peptide sequences were predicted using FragGeneScanPlusPlus^103^ commit 471fdf7 (fork: https://github.com/LeeBergstrand/FragGeneScanPlusPlus), via the ‘illumina_10’ model, from forward (R1) metagenome reads that passed the QC module of ATLAS (above). Peptide sequences were searched using the custom HMM developed in this study for “Ca. Chx. allophototropha”-associated PscA via hmmsearch 3.3.2^82^ with an e-value cutoff of 10^−10^. Raw hits were then filtered using a BLASTP^93^ search (BLAST version 2.10.1) against the PscA-like sequences of “Ca. Chx. allophototropha” and strain L227-5C. Only hits with >90% identity and with an e-value of <10^−10^ were retained. Predicted short peptide sequences were also searched using a HMM for the taxonomic marker gene rpoB from FunGene (June 2009 version)^104^ via hmmsearch 3.3.2^82^ with an e-value cutoff of 10^−10^. Counts of filtered PscA-like hits per metagenome were normalized to counts of RpoB hits following normalization by HMM length. Singleton filtered PscA-like hits were excluded from the analysis.

Metatranscriptome data were processed using the ATLAS pipeline^75^. The ‘qc’ module of ATLAS commit 59da38f was run to perform quality control of raw read data. Then, a customized fork of ATLAS, commit 96e47df (available under the ‘maprna’ branch at https://github.com/jmtsuji/atlas) was used to map RNA reads onto the set of dereplicated MAGs obtained from metagenome analyses (above) and to summarize RNA read counts. Briefly, the dereplicated MAGs were used as input for the “genomes” module of ATLAS so that QC-processed metatranscriptome reads were mapped onto the MAGs using BBMap. The minimum percent identity threshold for read mapping (‘contig_min_id’) was set to 99%. Following read mapping to MAGs, the counts of metatranscriptome read hits to genes within MAGs were summarized using featureCounts^105^, in version 1.6.4 of the Subread package. Default settings were used, except for the following flags: ‘-t CDS -g ID --donotsort’. Raw analysis code is available in the GitHub repository associated with this work.

After generating RNA read mapping data via ATLAS, the relative expression of each dereplicated MAG was calculated within each metatranscriptome. To perform this calculation, the number of RNA reads that mapped to each MAG was divided by the total number of RNA reads that mapped to all MAGs. The resulting relative expression values were averaged between replicate metatranscriptomes. In addition, we calculated expression levels of genes within RCI-encoding Chloroflexota MAGs based on normalization to dnaK and normalization by gene length, and we averaged gene expression levels between replicate metatranscriptomes (Supplementary Methods).

### Data availability

Enrichment culture metagenomes from the early to mid enrichment phase of the “Ca. Chx. allophototropha” culture (subcultures 1 and 15.2) and the L227-5C (primary enrichment) culture, along with recovered genome bins, are available under the National Center for Biotechnology Information (NCBI) BioProject accession PRJNA640240. Amplicon sequencing data will be made available under the same NCBI BioProject accession. The complete “Ca. Chx. allophototropha” genome, along with associated raw read and amplicon sequencing data, will be made available upon publication at BioProject accession PRJNA909349. Similarly, the complete Geothrix sp. L227-G1 genome and associated long read data (subculture 15.c) will be made available upon publication at BioProject accession PRJNA975665. Metagenome data from 2016, sequenced by the JGI, are available in the JGI Genome Portal under Proposal ID 502896. Environmental metagenome and metatranscriptome data from 2017-2018 are available under NCBI BioProject accession PRJNA664486. The full set of 756 metagenome-assembled genomes used for read mapping of metatranscriptome data are available in a Zenodo repository (doi:10.5281/zenodo.3930110) and will be made available on NCBI upon publication.

### Code availability

Custom scripts and additional raw data files used to analyze the metagenome and genome data are available at https://github.com/jmtsuji/Ca-Chlorohelix-allophototropha-RCI (doi:10.5281/zenodo.3932366).

---

**Extended Data Fig. 1.**
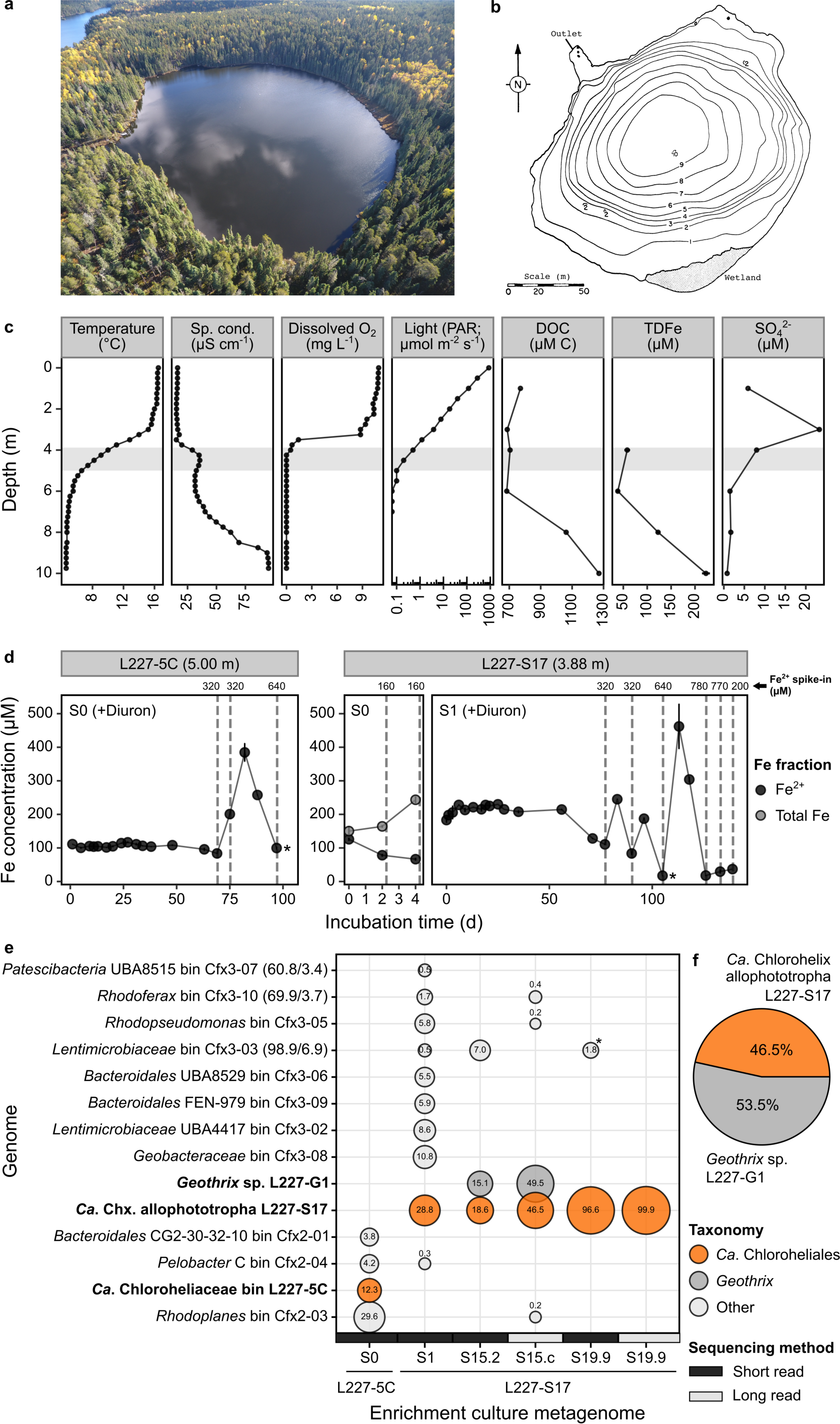
| Sampling and enrichment cultivation of “*Ca.* Chloroheliales” members. **a,** Aerial photograph of Lake 227 (IISD-ELA, Canada), facing eastward. **b**, Bathymetry map of Lake 227. Contour lines of 1 m are shown. **c**, Physicochemistry of the Lake 227 water column. A grey box marks the region of the water column from which water samples were collected for enrichment cultivation (top: 3.88 m; bottom: 5.00 m); these samples were collected during the same week as when samples for physicochemical parameters were collected. Standard deviations (n=2) are shown as error bars for TDFe measurements. **d**, Fe concentrations of the L227-5C and L227-S17 enrichment cultures over time. The primary enrichment (S0) is shown for the L227-5C culture, whereas the primary enrichment (S0) and first subculture (S1) are shown for the L227-S17 culture. Total Fe concentrations are only indicated for the L227-S17 primary enrichment, which was not spiked with Diuron. Asterisks mark the time points where samples were collected for metagenome sequencing. Standard deviations (n=2) of Fe concentrations are shown as error bars except for days 25, 28, and 139 of L227-S17 S1, where only single measurements were available. **e-f**, Microbial community composition the L227-5C and L227-S17 enrichment cultures over time. The bubble plot (**e**) shows the relative abundances of MAGs or closed genomes within enrichment culture metagenomes. Bubbles are shown where a genome recruited >0.1% of metagenome reads from a sample. The subculture generation of each enrichment is indicated by the number before the decimal place in its culture ID (e.g., S15.2 is a 15^th^ generation subculture). Estimated completeness and contamination (based on CheckM) are indicated in the taxon label for any MAG with <95% or >5% completeness or contamination, respectively. Nearly all (>95%) of the reads mapping to bin Cfx3-03 from the S19.9 short read metagenome (marked with an asterisk) were mapped to short contigs in Cfx3-03 that had >99% identity across 100% of their sequence to the “*Ca.* Chx. allophototropha” genome. The pie chart (**f**) shows the microbial community composition of a representative subculture (S21.2c) of the stabilized enrichment of “*Ca.* Chx. allophototropha” L227-S17. Relative abundances in the pie chart are based on 16S rRNA gene (V4 region) short read amplicon sequencing data with a detection sensitivity of 0.004%. Abbreviations: Sp. cond. Specific conductivity; PAR photosynthetically active radiation; DOC dissolved organic carbon; TDFe total dissolved iron. Aerial photograph and bathymetry map are courtesy of the IISD-ELA.

**Extended Data Fig. 2.**
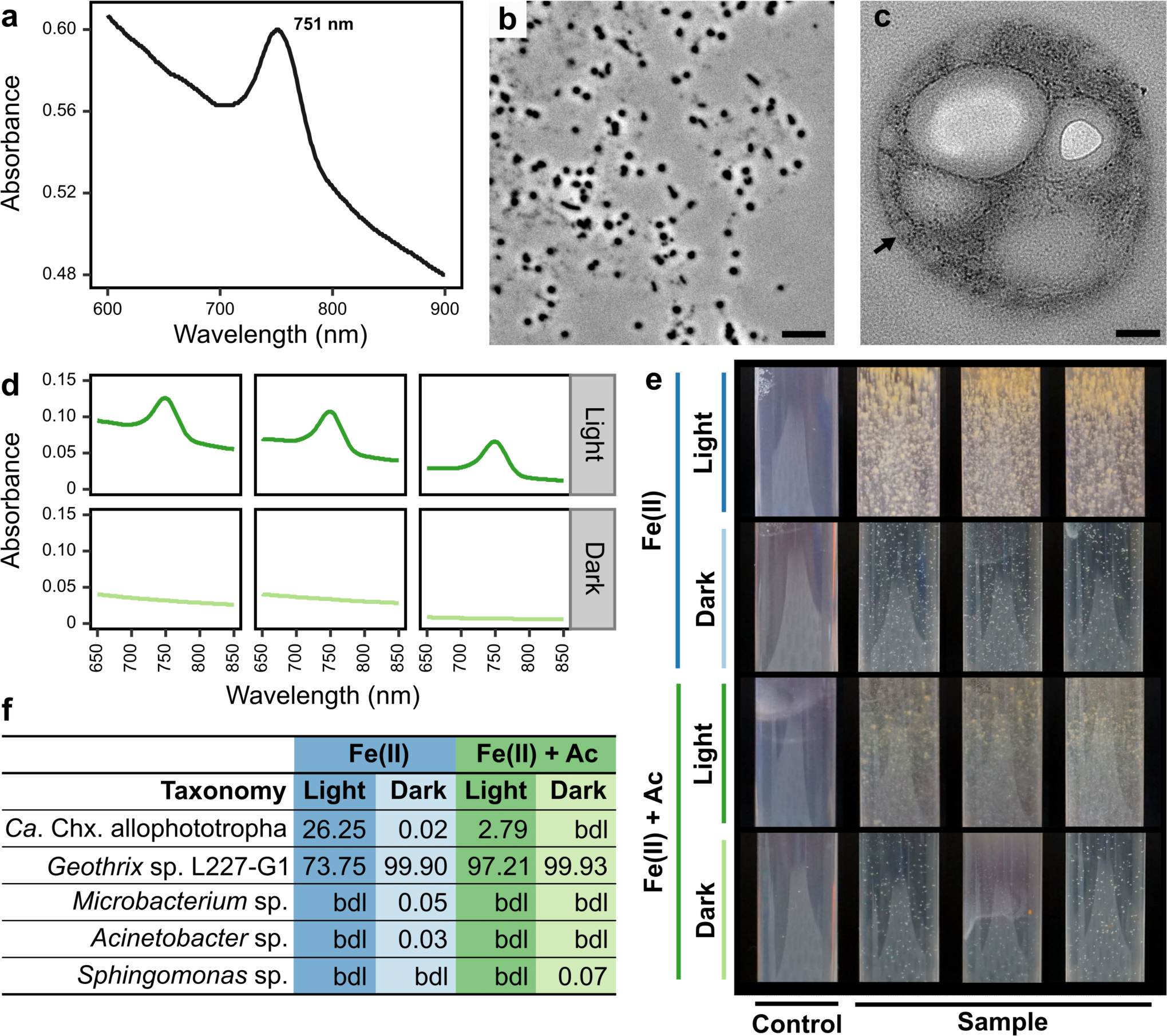
| Supporting data for the phototrophic physiology of “*Ca.* Chx. allophototropha”. **a**, Absorption spectrum of suspended whole cells from the “*Ca.* Chx. allophototropha” culture. The same sample was then sonicated before generating the *in vivo* spectrum in Fig. 1a. **b**, Light microscopy of *Geothrix* sp. L227-G1 (dry mount) grown under fermentative conditions. **c**, Transmission electron microscopy image showing a cross sections of “*Ca.* Chx. allophototropha” cells. An example chlorosome-like structure is marked with an arrow. **d**, Non-normalized absorption spectra for cultures grown in iron(II) with acetate in the light vs. dark; normalized data are shown in Fig. 1f. **e**, Appearance of agar shake tubes of “*Ca.* Chx. allophototropha” cultures (with resazurin) grown in the light or dark. Tube diameter is 1.55 cm. **f**, Microbial community of “*Ca.* Chx. allophototropha” cultures grown in the light or dark based on V1-V9 16S rRNA gene amplicon sequencing data. Data for iron(II) with acetate treatments are also shown in Fig. 1g but are shown here for comparison to iron(II) only (autotrophic) treatments. The scale bars in panels **b** and **c** represent 5 μm and 0.1 μm, respectively. Abbreviations: Fe(II) iron(II); Ac acetate.

**Extended Data Fig. 3.**
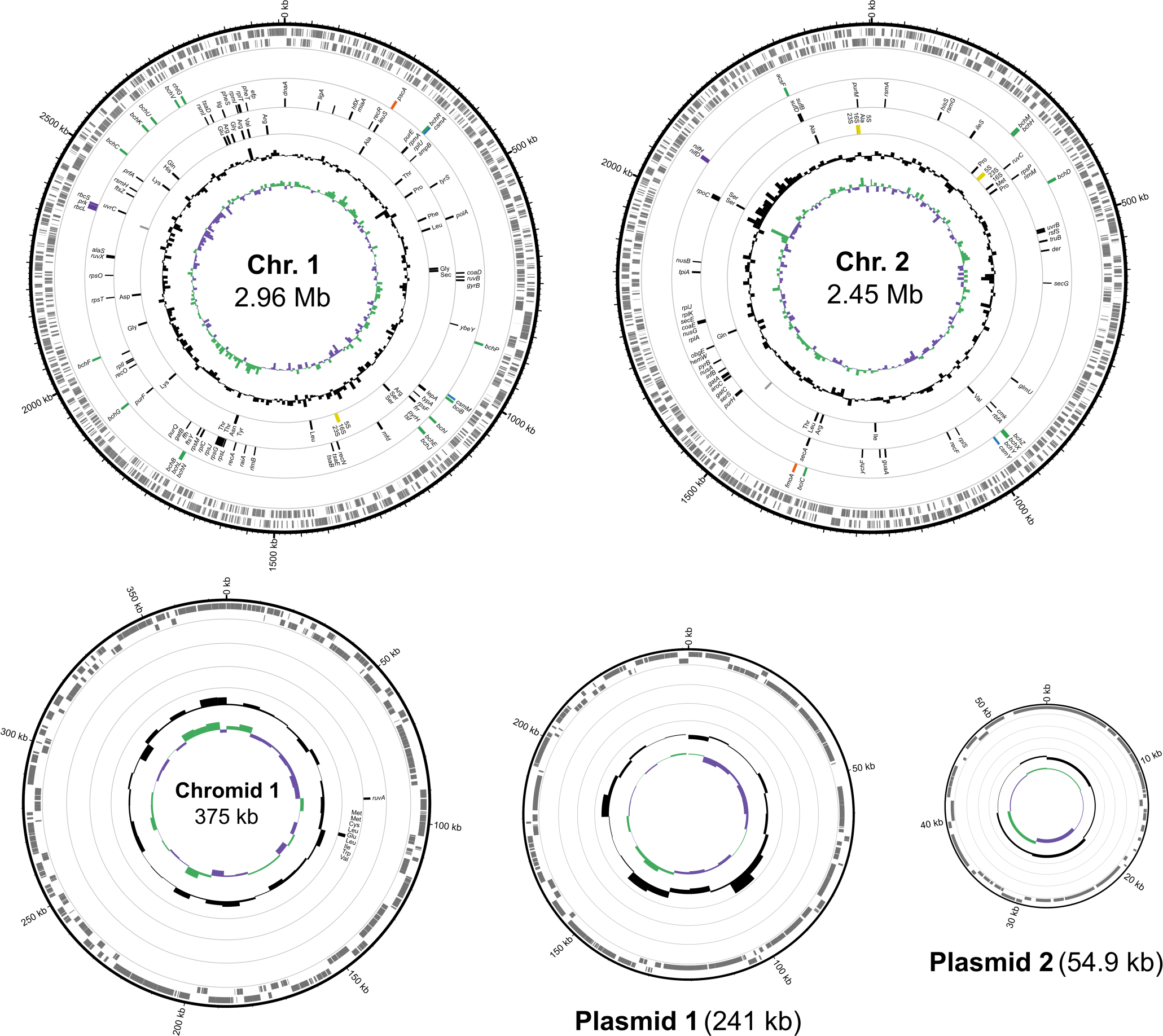
| Complete genome of “*Ca.* Chx. allophototropha”. A map of each of the five circular chromosomes in the “*Ca.* Chx. allophototropha” genome is shown. In each map, data rings (from outside to inside) show the following information: forward-orientation genes; reverse-orientation genes; photosynthesis-related genes as shown in Extended Data Table 2 (following the colouring scheme of Fig. 3); single copy marker genes identified by CheckM; non-coding genes including rRNA genes (yellow), tRNA genes (black), and others (grey); GC content, shown as ± 11.2% of the genome-wide average GC content of 46.8%; and GC skew, shown as ± 23.5%. The GC content and GC skew are shown based on a sliding window of 10 kb.

**Extended Data Fig. 4.**
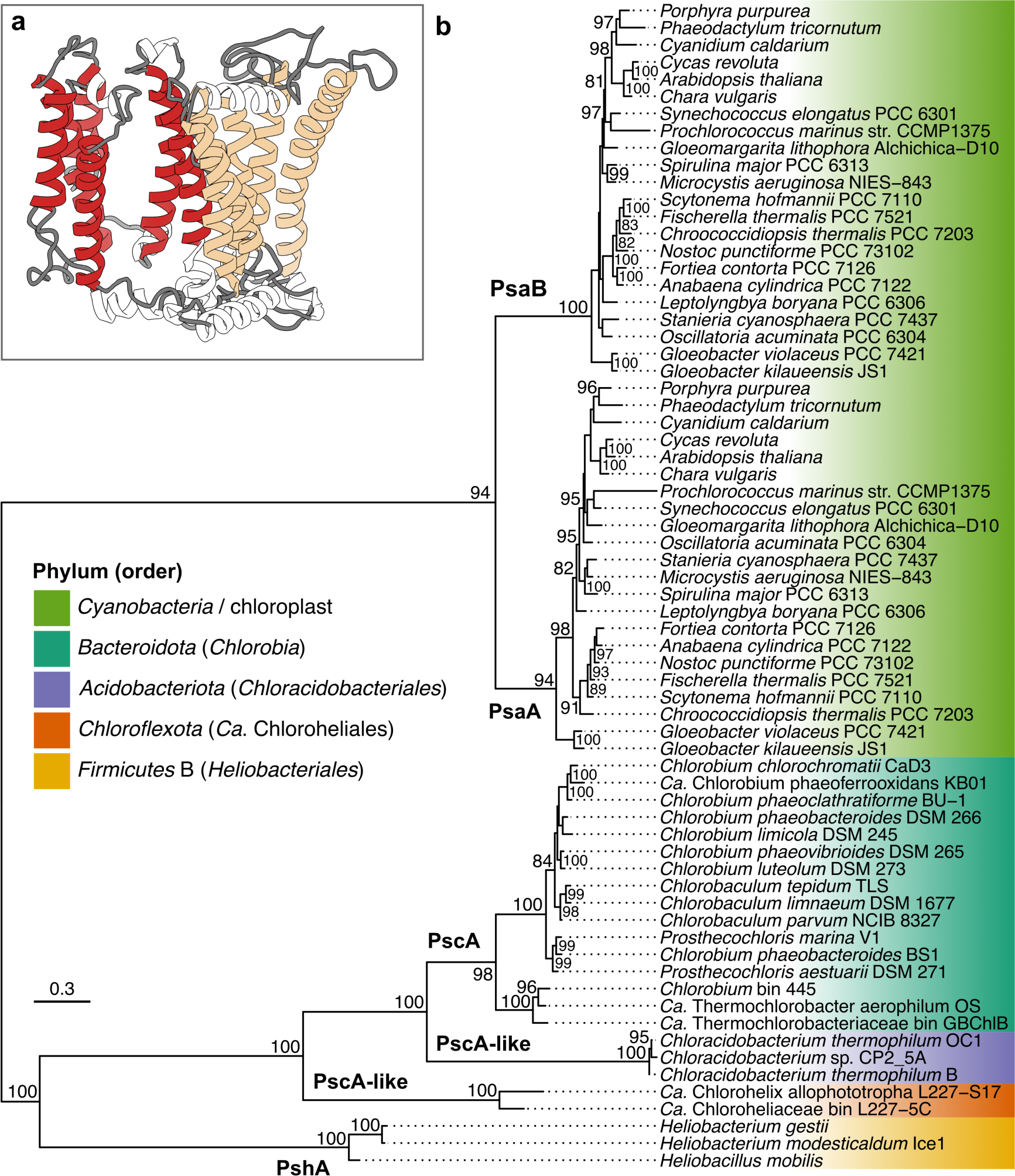
| Additional information about the Type I reaction center of “*Ca.* Chx. allophototropha”. **a,** Predicted tertiary structure of the novel “*Ca*. Chlorohelix allophototropha” PscA-like primary sequence based on homology modelling. The six N-terminal and five C-terminal transmembrane helices expected for RCI are coloured in red and tan, respectively. **b,** Maximum likelihood phylogeny of oxygenic and anoxygenic Type I reaction center predicted protein sequences. A simplified depiction of the same phylogeny is shown in Fig. 2. The phylogeny is midpoint rooted, and ultrafast bootstrap values of at least 80/100 are shown. The scale bar represents the expected proportion of amino acid change across the 548-residue masked sequence alignment.

**Extended Data Fig. 5.**
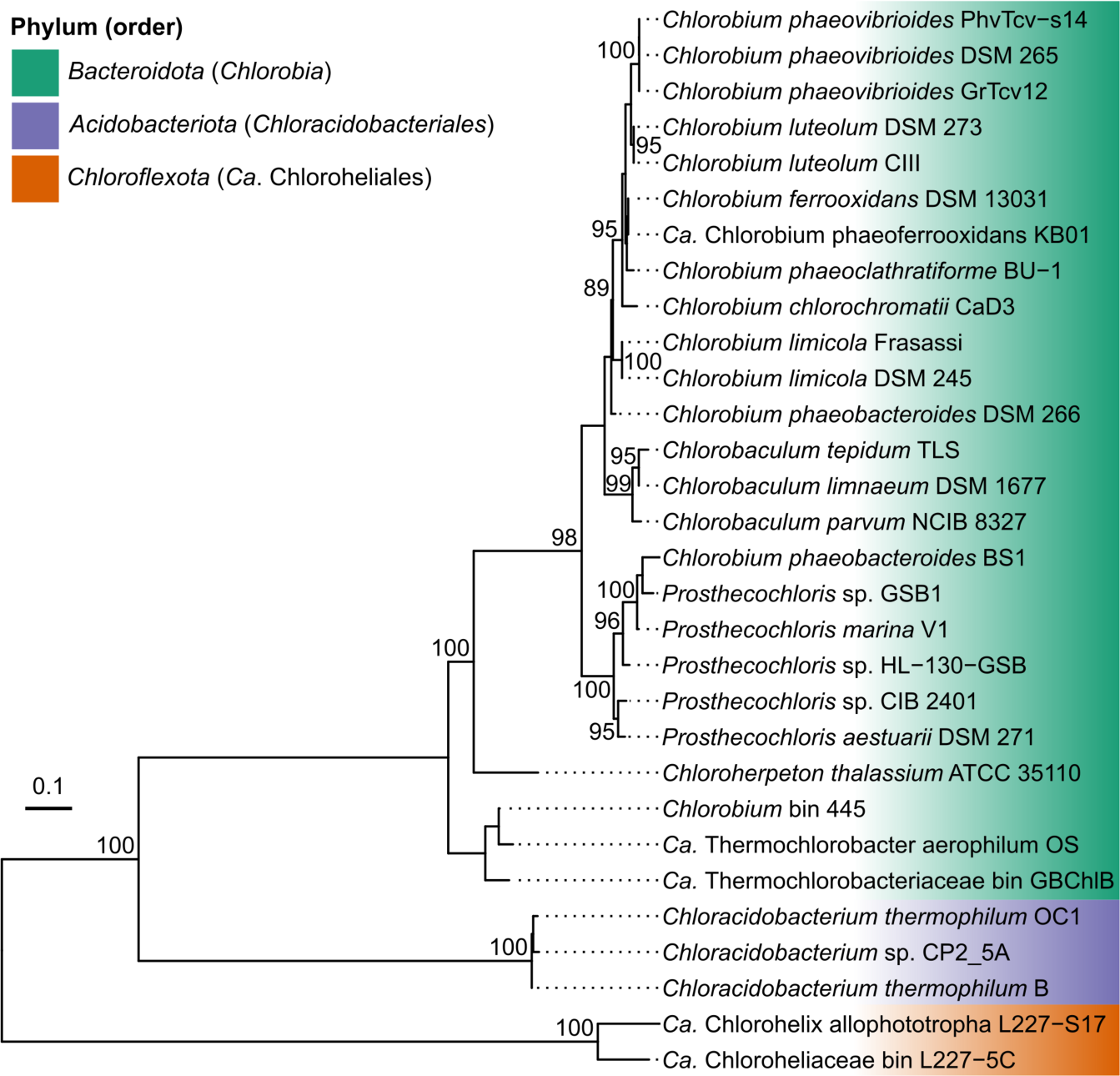
| Maximum likelihood phylogeny of Fenna-Matthews-Olson (FMO) protein (FmoA) sequences. The phylogeny is midpoint rooted, and ultrafast bootstrap values of at least 80/100 are shown. The scale bar represents the expected proportion of amino acid change across the 356-residue masked sequence alignment.

**Extended Data Fig. 6.**
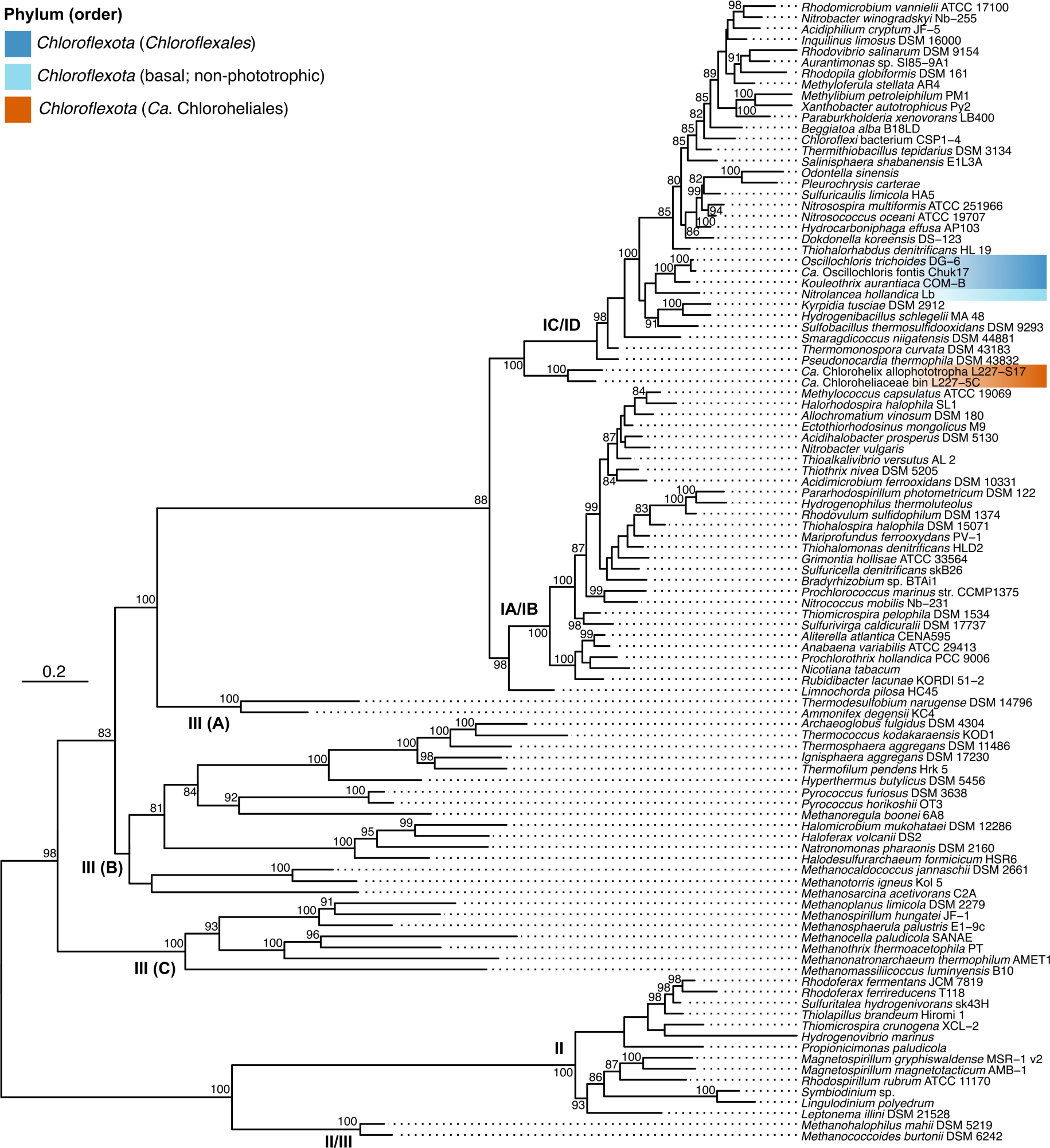
| Maximum likelihood phylogeny of RuBisCO large subunit (RbcL) predicted protein sequences. Group I to III RbcL sequences are shown, and group IV sequences, which do not form proteins involved in carbon fixation, are omitted for conciseness. The phylogeny is midpoint rooted, and ultrafast bootstrap values of at least 80/100 are shown. The scale bar represents the expected proportion of amino acid change across the 412-residue masked sequence alignment.

**Extended Data Fig. 7.**
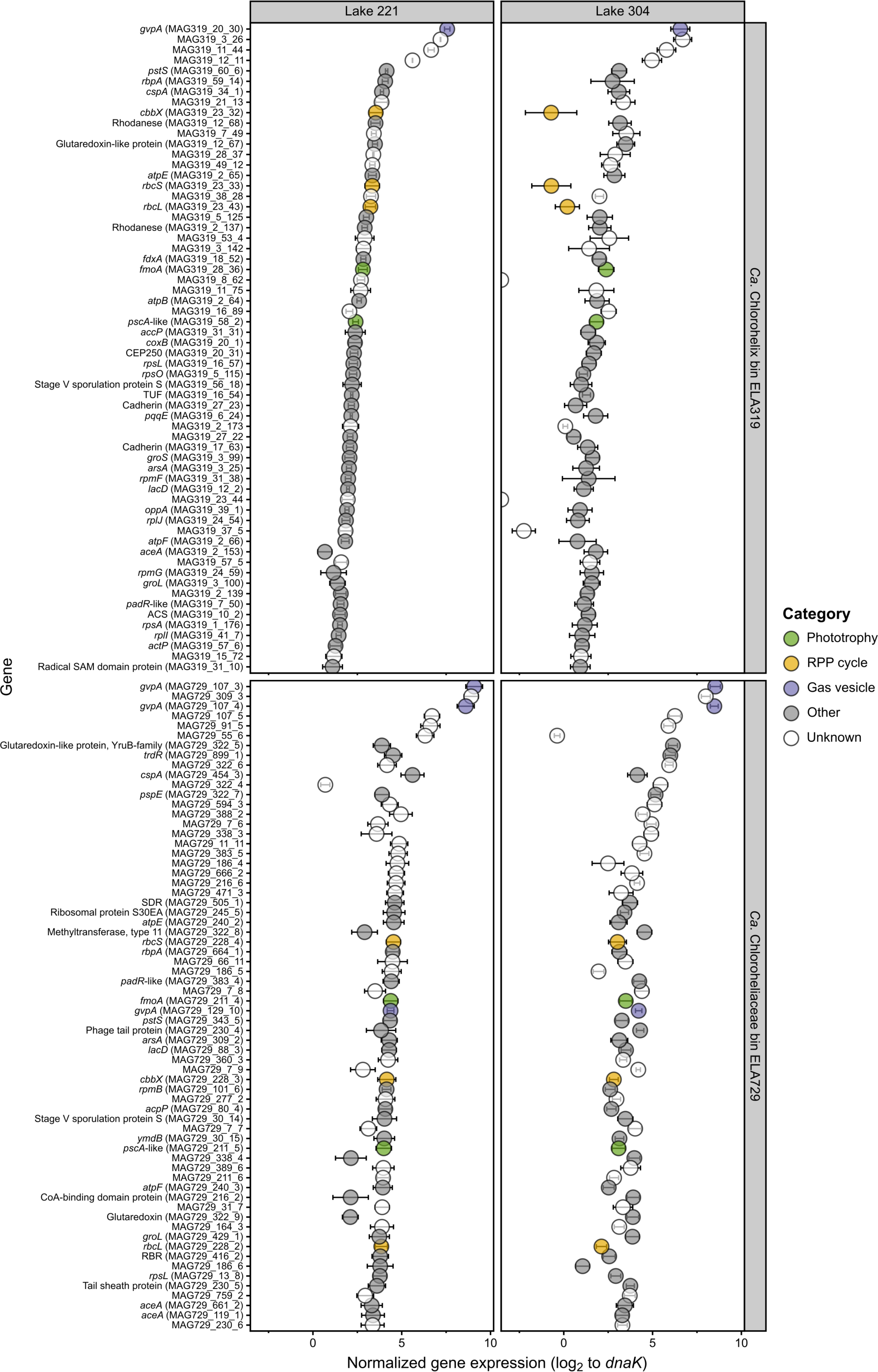
| Highly expressed genes among “*Ca.* Chloroheliales”-associated genome bins within Lake 221 and 304 metatranscriptome data. The top 50 protein-coding genes with highest normalized expression values are shown for genome bins ELA319 and 729. Error bars represent the standard deviation of the log_2_ expression ratio based on triplicate metatranscriptomes (n=3). Genes potentially involved in phototrophy, carbon fixation, and buoyancy are highlighted. Normalized expression values for all genes, along with their predicted amino acid sequences and annotations, are included in Supplementary Data 5.

**Extended Data Table 1.** | Microbial community composition of the “*Ca.* Chx. allophototropha” and *Geothrix* sp. L227-G1 enrichment cultures. Analyzed 16S rRNA gene amplicon data are shown to support spectroscopy, microscopy, and genomic analyses presented in this work. For each culture sample, the subculture generation is indicated (as part of the sample code), along with the figure panel(s) associated with that sample. Data was processed for each sequencing method as described in the methods. Sequences (i.e., ASVs or OTUs) were assigned as “*Ca.* Chx. allophototropha” or *Geothrix* L227-G1 based on >99% nucleotide match. Other ASVs or OTUs are presented in the “Other” category with their genus-level classification. Additional microbial community composition data, based on metagenome reads, are shown in Extended Data Fig. 1e. Abbreviations: bdl, below detection limit.

| Culture | Code* | Associated figure(s) | Sequencing method | Relative abundance (%) |  |  | Read count (detection limit, %)** | Notes |
| --- | --- | --- | --- | --- | --- | --- | --- | --- |
|  |  |  |  | “*Ca. Chx. allophototropa*” | *Geothrix* L227-G1 | Other |  |  |
| *Ca. Chlorohelix* | 13.2a-3 | 1d-e | Illumina V4-5 | 96.4 | 2.7 | 0.96 (*Dechloromonas*) | 523 (0.382) | 13.2a-3 was the parent culture of the culture used for SEM.<br>A parallel culture in the same agar dilution series as 13.2a-3 was used for TEM. |
| *Ca. Chlorohelix* | 19.9 | ED1e, ED3 | Nanopore V1-9 | 100.0 | Bdl | bdl | 28,321 (0.035) | Used for complete genome sequencing. |
| *Ca. Chlorohelix* | 21.2c | 1a-c, ED1f | Illumina V4 | 46.5 | 53.5 | bdl | 49,950 (0.004) | Two sequencing methods were compared for the same sample. |
|  |  |  | Nanopore V1-9 | 2.9 | 97.1 | bdl | 53,805 (0.019) |  |
| *Ca. Chlorohelix* | 22.2c | 1f-g, ED2e-f | Nanopore V1-9 | 2.8 | 97.2 | bdl | 65,971 (0.015) | Fe(II) + acetate, light |
| *Ca. Chlorohelix* | 22.2d | 1f-g, ED2e-f | Nanopore V1-9 | bdl | 99.9 | 0.07 (*Sphingomonas*) | 63,787 (0.016) | Fe(II) + acetate, dark |
| *Ca. Chlorohelix* | 22.2a | ED2e-f | Nanopore V1-9 | 26.3 | 73.8 | bdl | 50,792 (0.020) | Fe(II), light |
| *Ca. Chlorohelix* | 22.2b | ED2e-f | Nanopore V1-9 | 0.02 | 99.9 | 0.05 (*Microbacterium*),<br>0.03 (*Acinetobacter*) | 61,971 (0.016) | Fe(II), dark |
| *Geothrix* | 5.1b | ED2x | Nanopore V1-9 | bdl | 100.0 | bdl | 17,398 (0.057) | 5.1b was the parent culture of the culture used for microscopy. |

\*The number before the period (e.g., 12) refers to the subculture generation; subsequent numbers/letters were used internally to distinguish between cultures.
\*\*A sequence was detectable if it generated at least 2 counts (Illumina) or 10 counts (Nanopore).

**Extended Data Table 2.**
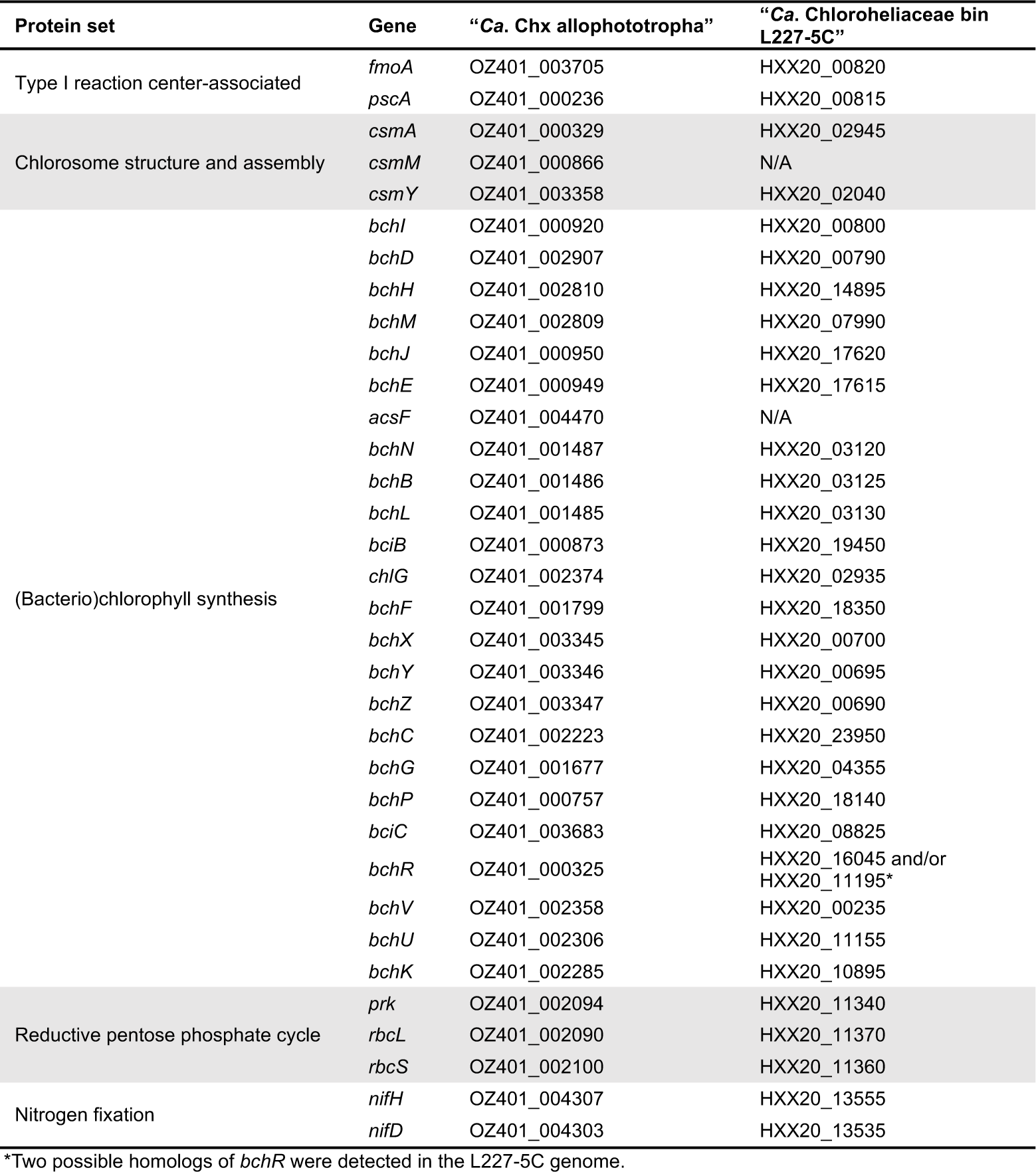
| Genes potentially involved in phototrophy or carbon/nitrogen fixation among genomes of “*Ca.* Chloroheliales” members recovered in this study. Locus tags are shown each gene. Results correspond to those shown in Fig. 3, except that homologs associated with the incomplete 3-hydroxypropionate bicycle are omitted for clarity, and additional genes involved in bacteriochlorophyll synthesis and the RPP cycle are shown.

**Extended Data Table 3.** | Summary of physicochemical parameters for sampled Boreal Shield lakes. The table includes the depth and surface area of all nine sampled lakes, along with information on the depths sampled for metagenome/metatranscriptome sequencing between 2016-2018. Summary parameters on the right side of the table show the topmost depth (sampled for metagenome sequencing) where dissolved oxygen was undetectable and the maximum measured concentrations of total dissolved iron, sulfate, and dissolved organic carbon among the collected samples. Full physicochemical data are provided in Supplementary Data 6. Abbreviations: n.d. no data.

| Lake | Maximum depth (m) | Surface area (m<sup>2</sup> x 10<sup>4</sup>) | Sampling year | Sampling month | Samples for metagenome sequencing (m) | Anoxic zone? (m) | Total dissolved iron (max.; µM) | Sulfate (max.; µM) | Dissolved organic carbon (max.; µM C) |
| --- | --- | --- | --- | --- | --- | --- | --- | --- | --- |
| L227 | 10.0 | 5.0 | 2016 | Jun | 6, 8, 10 | Yes (6) | 162.5 | 6.1 | 1092 |
|  |  |  | 2016 | Sep | 6, 8, 10 | Yes (6) | 188.1 | 8.6 | 1014 |
|  |  |  | 2017 | Sep | 1, 3, 4*, 6, 8, 10 | Yes (6) | 222.9 | 23.3 | 1271 |
| L221 | 5.5 | 9.0 | 2016 | Jun | 5 | Yes (5) | 15.2 | 11.7 | 844 |
|  |  |  | 2018 | Jul | 5* | Yes (5) | 68.4 | 12.2 | 922 |
| L304 | 6.0 | 3.6 | 2016 | Jun | 6 | Yes (6) | 37.6 | 9.3 | 676 |
|  |  |  | 2018 | Jul | 6* | Yes (6) | 137.6 | 8.2 | 865 |
| L222 | 6.0 | 16.4 | 2016 | Jun | 5 | No | 5.1 | 12.9 | 681 |
|  |  |  | 2016 | Sep | 5 | Yes (5) | 84.5 | 11.3 | 775 |
| L224 | 25.0 | 25.9 | 2016 | Jun | 25 | No | 0 | 17.8 | 244 |
|  |  |  | 2016 | Sep | 21, 25 | Yes (21) | 102.5 | 8.6 | 304 |
| L373 | 20.0 | 27.3 | 2016 | Sep | 20 | Yes (20) | 207 | 11.7 | 352 |
| L442 | 17.0 | 16.0 | 2016 | Jun | 9, 12, 15, 17 | Yes (15) | 170.7 | 18 | 759 |
|  |  |  | 2016 | Sep | 9, 13, 15, 17 | Yes (9) | 100.7 | 19.8 | 636 |
| L626 | 11.0 | 25.9 | 2016 | Jun | 11 | No | 0 | 12.8 | 361 |
|  |  |  | 2016 | Sep | 11 | Yes (11) | 19.6 | 15.1 | 363 |
| L239 | 30.0 | 54.3 | 2016 | Jun | 10, 20 | No | n.d. | 22.1 | 517 |
|  |  |  | 2016 | Sep | 10, 20 | No | 12.3 | 25.2 | 520 |
|  |  |  | 2018 | Jul | 20 | n.d. | n.d. | n.d. | n.d. |

\*Samples were also collected and used for metatranscriptome sequencing

## Supplementary Information | PDF file containing Supplementary Notes 1-3, Supplementary Methods, and Supplementary References

**Supplementary Data 1 | Amplicon sequencing variant table of early phototroph enrichment cultures from this study.** The Excel file summarizes the percent relative abundances of amplicon sequencing variants (ASVs) detected in 16S ribosomal RNA gene amplicon sequencing data representing early enrichment cultures of “*Ca.* Chloroheliales” members.

**Supplementary Data 2 | Bidirectional BLASTP results for photosynthesis-related genes among the *Chloroflexota* phylum.** The Excel file summarizes the query sequences and results of bidirectional BLASTP to search for photosynthesis-related genes in genomes of *Chloroflexota* members. These data are visualized in Fig. 3.

**Supplementary Data 3 | Mapping of metagenome reads to metagenome-assembled genomes.** The Excel file summarizes the percent mapped reads from each analyzed Boreal Shield lake metagenome to each of the 756 dereplicated MAG recovered in this study.

**Supplementary Data 4 | Mapping of metatranscriptome reads to metagenome-assembled genomes.** The Excel file summarizes the percent mapped reads from each analyzed Boreal Shield lake metatranscriptome to each of the 756 dereplicated MAG recovered in this study. For each metatranscriptome, the percentage of total reads that mapped to the MAG set is also summarized. Only MAGs with at least one mapped read are shown. Data for Lakes 221 and 304 are visualized in Fig. 4d.

**Supplementary Data 5 | Gene expression of “*Ca.* Chloroheliales”-associated genome bins.** The Excel file summarizes the normalized expression values of protein-coding genes in MAGs 319 and 729 based on environmental metatranscriptome data from Lakes 221 and 304. These data are visualized in Fig. 4e and Extended Data Fig. 7.

**Supplementary Data 6 | Physicochemical parameters for Boreal Shield lake samples.** The Excel file includes physicochemical parameters for all Boreal Shield lake water column samples used for metagenome/metatranscriptome sequencing in this work, except for the Lake 239 July 2018 sample, for which no physicochemical data are available. A summary of these data are presented in Extended Data Table 3. The file also includes light, temperature, and dissolved oxygen profile data from across the water columns of Lakes 221, 304, and 227 as visualized in Fig. 4c and Extended Data Fig. 1c.

