## Supplementary information for "Anoxygenic phototrophic *Chloroflexota* member uses a Type I reaction center"

This file contains Supplementary Notes 1-3, Supplementary Methods, and Supplementary References.

### Supplementary notes

*Supplementary note 1*

**Growth properties and metagenome-assembled genomes of enriched "*Ca.* Chloroheliales" members.** Liquid medium was used for the initial enrichments of strain L227-S17. However, filamentous phototrophs eventually ceased growth or were overtaken by purple phototrophic bacteria in subcultures made in liquid medium. Transfer of enrichment cultures to agar-containing medium allowed us to continue to culture strain L272-S17. As inoculum into agar-containing medium, we ultimately had to use a liquid enrichment (subculture generation 3) of the L227-S17 strain that had been stored at 4°C (dark) for five months, instead of an active culture that had been subcultured in liquid medium for additional generations, due to poor growth of the active culture. During growth in deep agar dilution series, additional selective conditions were used to purify the culture of specific contaminants. To eliminate purple phototrophic bacteria from the culture related to the metabolically versatile *Rhodopseudomonas palustris*[106], agar tubes were incubated under 740 nm LED lights, because light at ~740 nm wavelength is not readily used for photosynthesis by these bacteria[107]. Similar far-red LED lights were also used as a general light source for the "*Ca.* Chx. allophototropha" culture in later subcultures. The medium was adjusted to an elevated pH of ~7.5-8.5 for a time to select against green phototrophic bacteria belonging to the *Chlorobia* class, because these bacteria are known to grow poorly in moderately basic conditions[108]. Lastly, sodium molybdate was added to the medium at a final concentration of 100 µM, in a 1:10 concentration ratio relative to sulfate, to inhibit the activity of sulfate-reducing bacteria as optimized experimentally (Supplementary Methods). In addition, in some deep agar dilution series cultures, a carbonate-buffered sulfide feeding solution[57] was added at a final sulfide concentration of 100-120 µM to decrease the redox potential of the medium, although we did not observe an effect of this treatment on culture growth.

Physiologically, "*Ca.* Chx. allophototropha" had several shared characteristics with other cultured and RCII-utilizing phototrophs belonging to the *Chloroflexota* phylum. Cells grew well in soft agar and developed as long, spiralling filaments, similar to those observed for *Chloronema* spp., a group of poorly studied *Chloroflexota* members characterized primarily by microscopy from stratified lakes[109]. Although gliding motility of "*Ca.* Chx. allophototropha" cells was not observed, cells formed tight clumps, like *Chloroflexus aggregans*[18], that would readily bind to glass surfaces or to precipitates in culture medium. We were unable to grow "*Ca.* Chx. allophototropha" in freshwater medium amended with 0.5-3 mM sulfide in place of ferrous iron.

Although strain L227-5C was lost after the initial enrichment from L227, we recovered a high completeness but fragmented MAG of L227-5C from an enrichment culture metagenome. Following manual curation, the MAG was estimated by CheckM to have a completeness and contamination of 96.3% and 1.6%, respectively, and was composed of 546 contigs with a total length of 6.31 Mb and average GC content of 47.7 %. The MAG encoded 4794 genes, including 79 tRNA genes and a full-length 16S and 23S rRNA gene, and had a relative abundance of 12.3% within the enrichment culture metagenome based on recruitment of QC-processed short reads (Extended Data Fig. 1e). Three non-*Chloroflexota* genome bins were also recovered from the metagenome, including a RCII-encoding phototroph, classified to the *Rhodoplanes* genus, that likely corresponded to the purple phototrophic bacterium observed in the culture and had a relative abundance of 29.6% in the metagenome (Extended Data Fig. 1e). Genome bins classified to the *Pelobacter* genus and the CG2-30-32-10 family (in the *Bacteroidales* order), that had relative abundances of 4.2% and 3.8% in metagenome, respectively, were also recovered (Extended Data Fig. 1e).

*Supplementary note 2*

**Genomic potential for photosynthetic electron transport by "*Ca.* Chx. allophototropha".** We explored the potential photosynthetic electron transport chain used by "*Ca.* Chx. allophototropha" based on the strain's genomic sequence. Rather than detecting genes for Alternative Complex III, which is used by RCII-containing *Chloroflexota* members[39], we detected a potential *petXCDB* gene cluster (GenBank locus tags OZ401_002244 to OZ401_002247) homologous to the *petXDBC* gene cluster in *Heliobacterium modesticaldum* that encodes a cytochrome $b_6f$-like complex[40,110,111]. Unlike in *Chloracidobacterium thermophilum*, the *pscB* gene, encoding an iron-sulfur protein involved in electron transfer from RCI, did not occur in a cluster with *pscA* (see above). The "*Ca.* Chx. allophototropha" genome encoded numerous other iron-sulfur proteins that included the conserved eight cysteine residues observed in the PscB sequence of *Chlorobia* members and *Chloracidobacterium thermophilum*[9], and it is possible that one of these iron-sulfur proteins plays an analogous role in electron transfer in "*Ca.* Chx. allophototropha". In the same way, we identified multiple genes that could encode *c*-type cytochromes involved in electron transfer like *pscC* (associated with *Chlorobia* members)[28] or *petJ* (associated with *Heliobacteriales* members)[110]. We did not detect any putative homologs of the RCI-associated *pscD* gene encoded by *Chlorobia* members[28].

*Supplementary note 3*

**Diversity of the proposed "*Ca.* Chloroheliales" order.** Searching the GTDB[48] (release 89) revealed two MAGs associated with the proposed "*Ca.* Chloroheliales" (i.e., "54-19)" order that were assembled and binned in previous studies. One MAG, "*Chloroflexi* bin 54-19", was recovered from an ammonium sulfate bioreactor metagenome[112] and placed sister to the MAG of strain L227-5C that was recovered in this study (Fig. 3). The other MAG, "*Chloroflexi* RR_metagenome_bin16", was recovered from a polar surface soil metagenome (Robinson Ridge, Antarctica)[113] and placed basally to all other members of the "*Ca.* Chloroheliales" order (Fig. 3). Neither MAG contained detectable genes for chlorophototrophy (Fig. 3). Searching amino acid sequences predicted directly from the unassembled metagenome data used to generate these two bins, using the custom HMMs generated in this study (available in the code repository associated with this work), also did not reveal any RCI-related sequences other than those with a 100% match to chloroplast PscA (data not shown). These data suggest that not all members of the "*Ca.* Chloroheliales" are phototrophic.

**Supplementary methods**

*Additional details for the enrichment of novel Chloroflexota members*

Agar shake tubes initially used for selective enrichment of strain L227-S17 contained a freshwater medium[29] and 0.6% (w/v) triple-washed Bacto Agar (Becton, Dickinson and Company; New Jersey, U.S.A.). Agar plugs containing 10 mM ferrous chloride were added to the bottom of the tubes to form an iron concentration gradient, which was used to qualitatively determine favourable ferrous iron concentrations for growth. Medium was additionally amended with a 6 mM of an acetate feeding solution[57] to promote phototroph growth, 50 µL $L^{-1}$ of a supplementary vitamin solution[114], and 1.1 mM sodium thioglycolate as a reducing agent. Tubes were incubated at 18°C under halogen light (<50 µmol photons $m^{-2}$ $s^{-1}$). These conditions were gradually adjusted and optimized over subsequent subcultures to develop Chx3.1 medium, which is described in the Methods (and is described in more detail below).

The main contaminating bacterium in the "*Ca.* Chx. allophototropha" culture after multiple rounds of cultivation in deep agar dilution series, *Geothrix* sp. L227-G1, was selectively enriched to characterize the strain. Biomass was picked from an agar tube of "*Ca.* Chx. allophototropha" culture (subculture generation 21) and was inoculated into liquid freshwater medium[29] amended with 10 mM sodium acetate and 100 mM poorly crystalline iron(III) oxide[115], kept at pH 6.5-7.0. Ferrous chloride (1 mM) was also added as a reducing agent. Cultures were incubated in the dark at 22°C. Due to slow growth on poorly crystalline iron(III) oxide, the ferric iron source was changed in later subcultures to 2.8 mM ferric nitrilotriacetic acid. Reduction of ferric iron to ferrous iron was confirmed using the ferrozine assay[55]. To avoid precipitates in the medium, fermentative conditions were promoted in the culture using 10 mM trisodium citrate, with 0.25 mM ferrous chloride as a reducing agent, prior to microscopy. Culture samples for microscopy were washed once with Fe- and citrate-free medium (via centrifugation at 12,000 x g, 2 min), and a dry mount was then prepared and stained with crystal violet. Light microscopy (phase contrast) was performed using an Axioplan2 imaging microscope system equipped with an Axiocam MRm camera (Carl Zeiss; Oberkochen, Germany). Microscopy images were acquired using the AxioVision software, version 4.6.3 SP1 (Carl Zeiss).

*Details on Chx3.1 medium*

Chx3.1 medium had the following composition (chemical formulas and empirical units are included for reproducibility): $NH_4Cl$ (0.15 g $L^{-1}$), $MgSO_4 \cdot 7\ H_2O$ (0.25 g $L^{-1}$), $CaCl_2 \cdot 2H_2O$ (0.05 g $L^{-1}$), and optional resazurin (0.5 mg $L^{-1}$), which were autoclaved and cooled under $N_2$ gas. Sterile $KH_2PO_4$ and $NaHCO_3$ were then added to a final concentration of 0.3 g $L^{-1}$ and 0.93 g $L^{-1}$, respectively, along with 0.5 mL $L^{-1}$ trace element solution SLA (optionally selenite-free), 0.5 mL $L^{-1}$ of a previously published vitamin solution[56], 0.5 mL $L^{-1}$ selenite-tungstate solution[58], and 0.5 mL $L^{-1}$ of 50 mg $L^{-1}$ cobalamin. Concentrations of calcium pantothenate and thiamine were doubled (to 50 and 100 mg $L^{-1}$, respectively) in the vitamin solution compared to the reference. The pH was adjusted to 7.5 with stirring under 90:10 or 80:20 $N_2$:$CO_2$, and the medium was then dispensed into bottles that were stoppered with the same $N_2$:$CO_2$ headspace. This medium was amended with a final concentration of 0.2-0.8% molten agar, 2 mM ferrous chloride, and 1.2 mM acetate (either sodium acetate or an acetate feeding solution[57]) while preparing agar shake tubes, which were then kept stoppered under a $N_2$:$CO_2$

headspace. Triple-washed Bacto Agar (Becton, Dickinson and Company) or Agar A (Bio Basic; Markham, Canada) could be used as the agar source.

*Selection against sulfate-reducing bacteria in enrichment cultures*

Sulfate-reducing bacteria growing in enrichment cultures of "*Ca.* Chx. allophototropha" were enriched in Sulfate Reducing Medium (M803; HiMedia Laboratories; Mumbai, Maharashtra, India). Inoculum from the "*Ca.* Chx. allophototropha" culture was injected into Hungate tubes containing 10 mL of Sulfate Reducing Medium and a 90:10 $N_2$:$CO_2$ headspace, and tubes were incubated for seven days at room temperature in the dark. The enrichment was then subcultured into additional Hungate tubes containing Sulfate Reducing Medium that were spiked with sodium molybdate, a known inhibitor of sulfate reducing bacterial activity[116]. A molybdate concentration range of 0.2-200 mM was used, corresponding to a molybdate:sulfate ratio ranging from 1:100 to 10:1, given the 20 mM sulfate concentration in the medium. Tubes were incubated at room temperature in the dark for 14 days to monitor sulfate reducing bacterial activity, which was indicated by the development of a black iron-sulfide precipitate.

*Collection of biomass for physiological analysis*

Prior to spectroscopic characterization, cells from the "*Ca.* Chx. allophototropha" culture were picked and concentrated by centrifugation. Colonies were then picked out of the soft agar using a serological pipette, and picked material was centrifuged at 16-20,000 x g for 2 min at 4°C during each of three washes in phosphate buffered saline. During each wash, the golden portion of the pellet was selectively resuspended while discarding overlying agar and inorganic crystalline material that accumulated at the bottom of the pellet. Similarly, cells from the *Chlorobium ferrooxidans* culture were harvested and concentrated by centrifugation. Culture material was first centrifuged at 16,000 x g for 5 min at 4°C to collect a pellet containing cells mixed with abundant iron(III) oxides. This pellet was washed three times in phosphate-buffered saline, via centrifugation at 20,000 x g for 2 min at 4°C. After each wash step, the green layer that formed at the surface of the brown pellet was selectively resuspended for the subsequent wash, as described previously[101]. Lastly, cells from the *Chloroflexus aurantiacus* culture were harvested by centrifugation at 15,000 x g for 5 min at 4°C. The resulting pellet was washed 3 times in 10 mM Tris-Cl (pH=8) via centrifugation at 12,000 x g for 2 min at 4°C. Collected biomass was then used for spectroscopic analysis as described in the Methods.

*Electron microscopy*

To perform TEM analysis, cell pellets that had been digested with agarase and fixed in 4%/4% glutaraldehyde/paraformaldehyde were washed, enrobed in 4% Noble agar (ThermoFisher Scientific; Waltham, Massachusetts, USA), and cut into 1 mm cubes. The cubes were fixed in 1% (w/v) osmium tetroxide for 45 min and subjected to a 25-100% ethanol dehydration series. Samples were then subjected to stepwise infiltration using 25-100% LR White Resin (Ted Pella; Redding, California, USA), transferred to gelatin capsules in 100% LR White Resin, and allowed to polymerize overnight at 60°C. An Ultracut UCT ultramicrotome (Leica Microsystems; Wetzlar, Germany) equipped with a diamond knife (DiATOME; Hatfield, Pennsylvania, USA) was used to cut 50 nm thin sections, which were then floated on 100-mesh copper grids and stained with 2% uranyl acetate and Reynold's lead citrate to enhance contrast. Sections were imaged under standard operating conditions using a Tecnai G2 F20 TEM (ThermoFisher Scientific; Waltham, Massachusetts, USA) that was running at 200 kV and was equipped with a Gatan 4k CCD camera (Gatan; Pleasanton, California, USA).

For SEM analysis, fixed cells that had been agarase-treated were washed in phosphate-buffered saline and incubated in 1% (w/v) osmium tetroxide at room temperature for 30 min. Following incubation, cells were washed, deposited on an aluminum stub, dried, and then sputter coated with a gold:palladium mixture using a Desk V TSC Sample Preparation System (Denton Vacuum; Moorestown, New Jersey, USA). Prepared samples were imaged using a Quanta FEG 250 SEM (ThermoFisher Scientific) with a high-voltage setting of 10 kV and working distance of 9.9 mm.

*Long read amplicon sequence analysis*

NanoCLUST commit a09991c (fork: https://github.com/jmtsuji/nanoclust) was used to analyze long read amplicon sequencing data[74]. Default settings were used for analysis of post-demultiplexed samples, except Medaka (Oxford Nanopore Technologies) was upgraded to version 1.5.0 to use the 'r941_min_sup_g507' model for polishing, and the minimum cluster threshold was decreased to 10 reads. Following NanoCLUST, PCR primers and adaptors were trimmed off sequence clusters using CutAdapt version 3.4[73] in two rounds. In the first round, forward primers were removed, and the '--revcomp' flag was used to correct all sequences to the forward orientation. The reverse primer was removed in the second round. Any clusters without both forward and reverse primers were discarded. Trimmed sequences were then clustered at 99% identity[117] across all samples using the easy-cluster module (cov-mode 5, cluster-mode 2) of MMseqs 2 version 13.45111[118] to generate operational taxonomic units (OTUs). Chimera filtration was performed on clustered sequences using UCHIME2 version 11.0.667 (32 bit), using the NCBI 16S rRNA gene database for type strains (December 2022) as the reference. The known "*Ca.* Chx. allophototropha" 16S rRNA gene sequence (based on genome sequencing) was added to the NCBI database to improve chimera detection accuracy.

*Read cloud genome sequencing*

The "*Ca.* Chx. allophototropha" enrichment culture (subculture 15.2) was grown using white light (30 µmol photons $m^{-2}$ $s^{-1}$; mix of incandescent and fluorescent sources) with 5 mM acetate and an additional 100 µM molybdate. High molecular weight genomic DNA was then extracted from picked colony material using a modified salting-out procedure (Manual CG000116, Rev A; 10x Genomics; Pleasanton, California, USA), followed by a phenol:chloroform treatment modified from Zhou and colleagues[119]. Briefly, colonies were picked from agar, centrifuged at 1010 x g for 7 min, and washed once with phosphate-buffered saline to remove excess agar. A lysis buffer consisting of 1.33 mL of 10 mM Tris-HCl (pH=8), 400 mM NaCl, and 2 mM ethylenediaminetetraacetic acid (EDTA; pH=8) was added to the pelleted biomass. In addition, 0.53 mL of 10% sodium dodecyl sulfate and 0.5 mL of 1 mg/mL proteinase K (ThermoFisher Scientific; Waltham, Massachusetts, USA), 1% SDS, and 2 mM EDTA (pH=8) were added. Cells were then gently lysed for 18 hours at 37°C. Following lysis, 4.3 mL of 5 M NaCl was added, and the sample was centrifuged at 1010 x g for 5 min. The supernatant was transferred to 15 mL MaXtract High Density tubes (Qiagen; Venlo, The Netherlands), gently mixed with 1 volume phenol:chloroform:isoamyl alcohol (25:24:1), and centrifuged at 1500 x g for 2 min. The resulting 11 mL of supernatant was transferred into 19.5 mL of ice-cold 100% ethanol, mixed by inversion, and aliquoted into 1.5 mL tubes for ethanol precipitation. Tubes were centrifuged at 4°C and 6200 x g for 5 min to precipitate DNA. After removal of supernatant and drying of DNA pellets, the dry DNA pellets were serially re-suspended into a single 30 µL aliquot of Tris-EDTA (TE; 10 mM Tris-HCl and 1 mM EDTA; pH=8) buffer.

Size selection was performed on the resulting extract using pulsed-field gel electrophoresis and electroelution, as described previously[120]. Briefly, pulsed-field gel electrophoresis was performed using a CHEF MAPPER Pulsed-Field Gel Electrophoresis system (Bio-Rad; Hercules, California, USA) run at 14°C, 5.5 V/cm, 1.0-6.0 s pulse, and 120° angle, for 16 hours. Electroelution of DNA from the excised gel fragment, targeting DNA strands longer than 25 kb, was performed at 120 V for 2 h in cellulose membrane dialysis tubing (12,000 MWCO with 33 mm average flat width; Sigma-Aldrich; St. Louis, Missouri, USA), with current reversed for 1 min before completion. Dialysis tubing was prepared in advance of electroelution by boiling in a 2% sodium bicarbonate and 1 mM EDTA (pH=8) solution for 10 min, then boiling in water for 10 min, and finally storing in 20% ethanol and 1 mM EDTA (pH=8) at 4°C until use. Electroeluted DNA solution was concentrated using Amicon Ultra-15 Centrifugal Filter Units (30 kDa MWCO; Millipore; Burlington, Massachusetts, USA), and DNA was washed via ethanol precipitation and resuspended in 20 µL of TE buffer.

Library preparation for read cloud DNA sequencing was performed on the extracted DNA using the TELL-Seq WGS Library Prep Kit[121] (Universal Sequencing Technology; Canton, Massachusetts, USA) following ultralow input protocol recommendations for small genomes. Amplification of the library was performed using 10 µL TELL-beads and 16 amplification cycles. The library was sequenced using a MiSeq Reagent Kit v2 (300-cycle; Illumina) with 2x150 bp read length on a MiSeq System to a depth of 19.6 million total reads. Following sequencing, Picard version 2.21.6 (Picard Toolkit – Broad Institute – http://broadinstitute.github.io/picard/) was used to demultiplex output reads.

*Analysis of enrichment culture short read metagenome data*

Short read metagenome data generated for enrichment cultures of "*Ca.* Chx. allophototropha" L227-S17 and strain L227-5C were assembled and partitioned into MAGs to understand the microbial community compositions of the enrichments. For a metagenome of the early L227-5C enrichment (subculture 0), the ATLAS pipeline, version 2.2.0[75], was used for read quality control, metagenome assembly, and genome binning. A genome bin corresponding to strain L227-5C (based on classification via the GTDB-Tk[102] to the "54-19" order of the *Chloroflexota* phylum) was then manually curated (see next section) and was annotated using PGAP[85] during submission to the NCBI database. A metagenome of the early L227-S17 enrichment (subculture 1) and the read cloud metagenome of the more highly enriched version of the L227-S17 culture (subculture 15.2; described above) were also analyzed using ATLAS version 2.2.0. Read quality control and metagenome assembly were performed normally for the early enrichment (subculture 1) metagenome, but custom steps were needed to utilize the read cloud metagenome data (for subculture 15.2). Read quality control and metagenome assembly steps were first performed normally within the ATLAS pipeline for the read cloud metagenome. Then, to achieve a more contiguous assembly, quality control and assembly were performed again outside of ATLAS. The Tell-Read pipeline, version 0.9.7 (Universal Sequencing Technology), was used to re-analyze raw read outputs from the sequencer, performing demultiplexing and quality control on index reads via default settings, and the reads were then assembled using Tell-Link, version 1.0.0 (Universal Sequencing Technology), using global and local kmer lengths of 65 and 35, respectively. Following assembly using Tell-Link, the assembled scaffolds initially generated by ATLAS for the read cloud metagenome were substituted with the scaffolds generated using Tell-Link, and the ATLAS pipeline was then allowed to continue normally into the differential abundance genome binning step, which used MaxBin 2 version 2.2.4[99] and MetaBAT2 version 2.12.1[100]. (The same genome binning tools were used for analysis of the L227-5C sample.) All ATLAS settings for the analyses of the L227-5C and L227-S17 cultures are available in the code repository associated with this work.

*Manual genome bin curation*

To manually curate the genome bin of strain L227-5C, which was highly fragmented, all predicted protein sequences in the bin were taxonomically classified using the Kaiju webserver[122] with default settings. Any contigs having at least one hit classified within the *Chloroflexota* phylum were retained. Protein-coding sequences on other contigs were further screened using BLASTP[93] against the NCBI RefSeq database[123] (e-value cutoff = $10^{-10}$). Contigs having median coverage values greater than or less than one standard deviation of the mean contig coverage value across the genome bin were assessed gene-by-gene for signs of mis-binning. Other contigs having a BLASTP hit with >80% identity to a RefSeq entry or containing a tRNA or rRNA gene (which was subsequently screened using BLASTN against the NCBI nr database, excluding uncultured subjects) were also examined.

In addition, although we eventually closed the "*Ca.* Chx. allophototropha" genome, we curated an earlier MAG of "*Ca.* Chx. allophototropha", which was recovered from read cloud metagenome sequencing data (as described in the above section), for preliminary study of the strain. To manually curate the MAG, genes predicted by Prodigal version 2.6.3[83] were queried against the NCBI RefSeq protein database[123] using BLASTP[93] (e-value cutoff = $10^{-10}$), and the taxonomic lineages of gene hits was determined using the script *make-lineage-csv.py* (https://github.com/dib-lab/2018-ncbi-lineages, commit 63e8dc7). Any scaffolds with at least one hit to a subject sequence associated with the phylum *Chloroflexota* were kept, and the top five highest-scoring hits for each gene were considered. Remaining scaffolds were checked for their median coverage, length, and gene classifications, and genes were further assessed against the NCBI nr protein database via BLASTP. Short contigs containing no genes or genes consistently matching the same non-*Chloroflexota* phylum were discarded from the bin. This curated MAG is available as NCBI accession GCA_013390565.1 but was not used for the genomic analyses in this work.

*Genome sequencing of Geothrix sp.* L227-G1

In addition to read cloud sequencing (of subculture 15.2), long read sequencing was additionally performed to close the genome bin of *Geothrix* sp. L227-G1. From a separate culture (subculture 15.c), cells were grown using 0.5-0.6% (w/v) agar, with the same light source and additional molybdate as subculture 15.2 (i.e., 30 µmol photons $m^{-2}$ $s^{-1}$ white light, mix of incandescent and fluorescent sources; 100 µM molybdate), and were harvested via centrifugation of the whole agar slurry at 20,000 x g for ≥30 min at 4°C. The resulting pellets were then used for a total of eight DNA extractions using the DNeasy UltraClean Microbial Kit (Qiagen), and output DNA was concentrated using the DNA Clean and Concentrator-5 kit (Zymo Research). A DNA sequencing library was then prepared via the Ligation Sequencing Kit (SQK-LSK110; Oxford Nanopore Technologies) with Long Fragment Buffer, using

370 ng of the concentrated DNA, combined with a spike-in of 125 ng of Lambda DNA (EXP-CTL001; Oxford Nanopore Technologies), as input. The library was sequenced using a R9.4.1 Flongle flow cell (FLO-FLG001; Oxford Nanopore Technologies). Adaptive sampling was used to deplete reads matching the Lambda phage genome (NC_001416.1) during sequencing. Basecalling was performed using Guppy 5.0.16 (Oxford Nanopore Technologies) with the Super Accuracy model, generating 0.23 million reads with a mean length of 3.9 kb.

Following sequencing, we combined long read data from subculture 15.c with short read (read cloud) data from subculture 15.2 to obtain a closed genome bin *Geothrix* sp. L227-G1. We obtained a set of assembled and polished contigs from the hybrid short- and long-read data using Rotary commit fd5acee (workflow is described in the Methods). Quality control of short reads was performed using ATLAS 2.2.0, using reads demultiplexed with Picard, prior to running Rotary. We then mapped the short read data (i.e., from subculture 15.2) to the contigs using BBMap version 37.99 (Bushnell B.), using a minimum sequence identity threshold of 95%, and performed genome binning using MetaBAT2 version 2.15[100]. This analysis resulted in a single-contig, closed and circular genome bin that had 16S rRNA gene sequences with >99% match to *Geothrix* sp. L227-G1 16S rRNA gene amplicon sequences. All other contigs in the assembly were associated with "*Ca.* Chx. allophototropha", except an 8.1 kb linear contig that had 10-fold lower coverage depth than any chromosomal sequence. We annotated the closed circular genome bin using PGAP version 2022-04-14.build6021[85].

*Microbial community composition of enrichment cultures*

To compare the microbial community composition of enrichment cultures across all analyzed samples, we clustered MAGs and genomes constructed from the L227-5C metagenome (subculture 0), L227-S17 short read metagenomes (subcultures 1 and 15.2), and the closed genome assemblies of "*Ca.* Chx. allophototropha" and *Geothrix* sp. L227-G1, using FastANI version 1.33[124] with a clustering threshold of 97.5%. When more than one genome belonged to a cluster, which only occurred in the case of "*Ca.* Chx. allophototropha" and *Geothrix* sp. L227-G1, we used the closed circular genome version as the cluster representative. All short read metagenome data were processed using the 'qc' module of ATLAS 2.8.2[75]. We then mapped the QC-processed short-read metagenome data to the clustered genome set using BBMap version 37.99 (Bushnell B.), using a minimum percent identity threshold of 90% and the 'ambiguous=best' setting. Similarly, we mapped all QC-processed long-read metagenome data (QC performed using Rotary, above) to the clustered genome set using Minimap2 version 2.23[125], and we excluded secondary or supplementary alignments using samtools 1.15[126]. Relative abundances were calculated based on the percent recruitment of reads to genomes.

*Collection of genomes/genes for phylogenomics*

To compare the genomic properties of *Chloroflexota* phylum members, representative genomes associated with the phylum were downloaded from NCBI based on information in GTDB[48] release 89. All genomes that represented a type species within the *Chloroflexota* according to NCBI and the GTDB were downloaded, as well as genomes representing members of known phototrophic clades (i.e., the *Chloroflexaceae* family[24], the "*Ca.* Thermofonsia" order[25], and the "*Ca.* Roseiliniales" order[19]). Selected genomes of known phototrophs that were deposited in other genome databases were also downloaded (i.e., genomes of "*Ca.* Chloranaerofilum corporosum"[19], "*Ca.* Roseilinea gracile"[19], and "*Ca*. Chlorothrix halophila"[21]; see details in the code repository associated with this work). In addition, any genome bins listed in the GTDB that belonged to the "54-19" order (which represented the name of the "*Ca.* Chloroheliales" order in this database) were downloaded. Non-phototrophic lineages were subsequently pruned to one representative per genus, except for non-phototrophic lineages closely related to phototrophic clades. In total, this left 58 genomes, including genomes of 28 known phototrophs. This genome collection was used for subsequent creation of a *Chloroflexota* species tree, as described in the Methods.

We collected additional genomes to compare photosynthesis genes encoded by "*Ca.* Chloroheliales" members to genes of other phototrophs. Genomes containing homologs of the genes of interest (*pscA*/*pshA*/*psaAB*, *fmoA*, *csmA*, *bchIDH*/*chlIDH*, *bchLNB*/*chlLNB*, *bchXYZ*, and/or *rbcL*; see Methods) were determined using a combination of automated detection via AnnoTree[127] and descriptions in the literature. Representative genomes from this initial genome set were selected from the GTDB, based on genome quality and taxonomic novelty, before being downloaded from NCBI. Where needed, genome nucleotide files were annotated using Prodigal 2.6.3[83]. Potential orthologs of interest were then identified in the downloaded genomes using bidirectional BLASTP, which was performed using the primary sequences of known reference genes as queries (shown in Supplementary Data 2) via BackBLAST[94] version 2.0.0-alpha3. In addition, for some sequence sets (i.e., Type I reaction centers; Bch proteins; RbcL), additional reference sequences were added manually based on literature references[28,95,96]. Sequence sets were then used to build photosynthesis gene phylogenies as described in the Methods.

*Searching for additional RCI and RCI-associated gene homologues*

Preparation of the collection of *Chloroflexota*-associated genomes revealed that two existing genomes bins in the GTDB placed taxonomically within the "*Ca.* Chloroheliales" order. Unassembled read files for the environmental metagenomes associated with those bins were downloaded from the European Nucleotide Archive (ENA) to identify any potentially unbinned but novel photosynthesis-associated genes. Short protein sequences were predicted directly from unassembled read data using FragGeneScanPlusPlus[103] commit 9a203d8. These short protein sequences were scanned using the two custom HMMs developed in this study (above) via hmmsearch[82] v3.1b2 using a relaxed e-value cutoff of $10^{-1}$. The average coverage and read recruitment of each genome bin was also calculated by mapping the unassembled metagenomic reads onto the bins using bbmap.sh version 38.75 (BBMap – Bushnell B.).

*Lake sampling events*

We sampled the water columns of eight seasonally anoxic lakes (Lakes 221, 222, 224, 227, 304, 373, 442, and 626) within the IISD-ELA, along with a permanently oxic reference lake (Lake 239), for DNA and/or RNA across four main sampling events. Sampling in June 2016, September 2016, and September 2017 involved multi-depth hypolimnion profiling of Lakes 227 and 442, for which metagenome data have been reported previously[101], along with mid-hypolimnion sampling of other selected lakes. A full water column profile of DNA samples was collected for Lake 227 in September 2017, along with collection of RNA samples from a single depth of Lake 227 in the upper anoxic hypolimnion. Lastly, in July 2018, Lakes 221 and 304 were surveyed again. A single depth in the mid anoxic hypolimnion of both lakes was sampled for both DNA and RNA.

*Detailed procedures for environmental DNA/RNA extraction*

For each Sterivex filter used for DNA collection, the Sterivex filter case was opened, and the filter membrane was carefully removed using a flame-sterilized scalpel blade. Each filter membrane was cut in half lengthwise along the filter. One half was stored frozen to provide a backup sample, and DNA was extracted from the other half using the DNeasy PowerSoil or DNeasy PowerSoil HTP 96 Kit (Qiagen; Venlo, The Netherlands). Extractions were performed according to the kit protocol, and the optional 10 min incubation at 70°C after adding Solution C1 was performed to enhance cell lysis. Mechanical lysis was performed for samples in PowerBead Tubes using a FastPrep-24 instrument (MP Biomedicals; Santa Ana, California, U.S.A.) set at 5 m/s for 45 s, and mechanical lysis was performed for samples in PowerBead DNA Plates using a mixer mill MM 400 (Retsch; Haan, Germany) set at 30 Hz for 10 min. Resulting DNA concentrations were then quantified using a Nanodrop spectrophotometer (Thermo Fisher Scientific) or using the Qubit dsDNA HS Assay Kit with Qubit 2.0 fluorometer (Thermo Fisher Scientific).

RNA extraction was performed using the ZymoBIOMICS DNA/RNA Miniprep Kit (Zymo Research) with initial steps of the protocol modified slightly to accommodate the volume of DNA/RNA Shield associated with each filter. Sterivex filters filled with DNA/RNA Shield were thawed, and the DNA/RNA Shield was pumped out of the filter cases and saved for downstream use. Filter cases were opened and filters excised as described above. Each filter half was cut into small pieces. One half was stored in a clean 2 mL microfuge tube along with half of the collected DNA/RNA Shield solution and frozen as a backup sample. The other filter half, along with the remaining DNA/RNA Shield, was added into a dry ZR BashingBead Lysis Tube for extraction. Mechanical lysis was performed using a FastPrep-24 instrument (MP Biomedicals). Lysis tubes were shaken at 6.5 m/s for 60 s twice, and tubes were allowed to cool on ice for at least 60 s between mechanical lysis rounds. After centrifugation, the entire supernatant volume was transferred to a new microfuge tube, and one volume of DNA/RNA Lysis Buffer was added to the tube and mixed. RNA extraction was then performed according to the "DNA & RNA Parallel Purification" protocol in the kit manual using in-column DNase I treatment. Only RNA (and not DNA) extracts were saved because corresponding Sterivex filters for DNA were collected and processed for the same lake depths using the protocol described above. The resulting RNA extracts were quantified using a Nanodrop spectrophotometer (Thermo Fisher Scientific) and Qubit RNA Assay Kit with Qubit 2.0 fluorometer (Thermo Fisher Scientific). Extracts were also run on a 1% agarose gel stained with GelRed (Biotium; Fremont, California, U.S.A.) to confirm that rRNA of the expected lengths was visible.

*Gene expression calculations*

To calculate gene expression levels based on environmental metatranscriptome data, the single-copy taxonomic marker gene *dnaK* was identified in each RCI-encoding *Chloroflexota* MAG that was recovered from Boreal Shield lake metagenome data. Homologs of *dnaK* were identified based on eggNOG-mapper[128] annotations (version 1.0.3), and annotations were confirmed using a BLASTP[93] search of the predicted protein sequence against the RefSeq[123] database. In the case of bin ELA729, two genes were annotated as *dnaK*, but one of the genes had low predicted sequence identity at the amino acid level to DnaK proteins in RefSeq (~34% identity) and most closely matched DnaK encoded by members of *Chitinophaga* spp. in the *Bacteroidetes* phylum, so this gene was not used for normalization. Relative expression levels of each gene within a MAG were then calculated by dividing the gene length-normalized hit count of each gene by the gene-length normalized hit count of *dnaK*, and expression levels were averaged among replicate metatranscriptomes. Functional annotations of genes with high relative expression were determined by eggNOG-mapper as part of the ATLAS 'Genecatalog' module, run on metagenome data, combined with manual annotation of photosynthesis-associated genes.

*Geospatial data processing*

A map of lakes at the IISD-ELA was generated using major and minor water body information from the CanMap Water dataset (DMTI Spatial; Markham, Canada). Topographical data from a digital elevation model (30 m intervals; DMTI Spatial) was displayed as 20 m contour lines on the map. In addition, the approximate range of Boreal Shield regions on Earth was determined using a subset of the Global GIS geospatial data collection (Esri; Redlands, U.S.A.). Regions defined as the major habitat "Boreal forest/taigas" (within the World Wildlife Fund Ecoregions dataset; wwf_eco.shp), were intersected with geologic provinces that included the term "Shield" in their entry name (within the

Geologic Provinces of the World dataset, United States Geological Survey; wep_prvg.shp). Intersection calculations were performed using ArcGIS (Esri). The resulting data were visualized on a Natural Earth basemap consisting of 1:50 m land, 1:50 m ocean, and 1:110 m lake vectors. All map visualizations were performed using QGIS (QGIS Association).
